# Global H-NS counter-silencing by LuxR activates quorum sensing gene expression

**DOI:** 10.1101/738146

**Authors:** Ryan R. Chaparian, Minh L. N. Tran, Laura C. Miller Conrad, Douglas B. Rusch, Julia C. van Kessel

**Affiliations:** Department of Biology, Indiana University, Bloomington, IN; Department of Chemistry, San Jose State University, San Jose, CA; Center for Genomics and Bioinformatics, Indiana University, Bloomington, IN

**Keywords:** LuxR, vibrio, quorum sensing, H-NS

## Abstract

Bacteria coordinate cellular behaviors using a cell-cell communication system termed quorum sensing. In *Vibrio harveyi*, the master quorum sensing transcriptional factor LuxR directly regulates >100 genes in response to changes in population density. Here, we show that LuxR derepresses quorum sensing loci by competing with H-NS, a global transcriptional repressor that oligomerizes on DNA to form filaments and bridges. We first identified H-NS as a repressor of bioluminescence gene expression, for which LuxR is a required activator. In an *hns* deletion strain, LuxR is no longer necessary for transcription activation of the bioluminescence genes, suggesting that the primary role of LuxR is to displace H-NS to derepress gene expression. Using RNA-seq and ChIP-seq, we determined that H-NS and LuxR co-regulate and co-occupy 28 promoters driving expression of 63 genes across the genome. ChIP-PCR assays show that as autoinducer concentration increases, LuxR protein accumulates at co-occupied promoters while H-NS protein disperses. LuxR is sufficient to evict H-NS from promoter DNA *in vitro*, which is dependent on LuxR DNA binding activity. From these findings, we propose a model in which LuxR serves as a counter-silencer at H-NS-repressed quorum sensing loci by disrupting H-NS nucleoprotein complexes that block transcription.

## Introduction

In nature, bacteria live in complex communities of microbes where competition for resources is constant. Thus, detection and identification of neighboring cells in the community is important for survival in many niches. To achieve this feat, bacteria use cell-cell communication called quorum sensing (QS) to detect and differentiate both the number and type of cells in the local vicinity (1). QS employs small signaling molecules called autoinducers (AIs). Because AIs are constitutively produced and released, their local extracellular concentration functions as a proxy for the number of neighboring cells. The marine pathogen *Vibrio harveyi* produces three AIs: HAI-1 (Harveyi autoinducer 1), CAI-1 (Cholerae autoinducer 1), and AI-2 (autoinducer 2) (reviewed in (2, 3)). Each of these AI molecules are sensed and bound by a cognate membrane-bound receptor: HAI-1 is detected by LuxN, CAI-1 is detected by CqsS, and AI-2 is detected by LuxPQ. At low cell density (LCD), when the cellular concentration of a population is low, AI concentration is relatively low, and the receptors remain unbound and thereby function as kinases. The phosphorylation cascade and downstream regulators converge to control the production of the master QS transcription factors, AphA and LuxR. At LCD, AphA is maximally produced and LuxR is expressed at its lowest level (4). As the population grows and transitions to high cell density (HCD), the AI concentration exceeds a threshold in which the receptor proteins are saturated by AI molecules. In the ligand-bound state, the receptor proteins change from kinases to phosphatases, switching the flow of phosphate and altering the expression and/or activity of downstream regulators. The end result is that, at HCD, LuxR is produced maximally, and AphA protein production is inhibited. This regulatory network results in the activation and repression of hundreds of genes in response to changes in population density (5, 6). The core of the QS signal transduction network architecture and the LuxR global regulator are conserved in *Vibrio* species, though the signaling molecules and/or receptors vary (6). Thus, in response to increases in population density and accumulating AIs, *Vibrio* cells increase production of LuxR protein, which results in a corresponding change in gene expression and behavior (*e.g.*, bioluminescence, competence, and secretion of virulence factors).

LuxR is a global regulator that controls the expression of >400 genes (5–8). This family of LuxR-type proteins is conserved across vibrios (*e.g.*, HapR in *Vibrio cholerae*, SmcR in *Vibrio vulnificus*), but does not bind AI as in the LuxI/LuxR systems widely found in Gram-negative bacteria. Rather, LuxR is classified as a TetR-type transcriptional regulator and can activate as well as repress gene expression. Our previous study showed that LuxR functions synergistically alongside the nucleoid-organizing protein called integration host factor (IHF) to activate transcription of the bioluminescence operon *luxCDABE* (7). Another study from our lab showed that LuxR interacts directly with the alpha subunit of RNA polymerase (RNAP) and that this interaction is required for activation of a subset of QS genes (9). These findings suggest that LuxR-dependent transcriptional activation requires the use of accessory proteins to remodel DNA structure and position RNAP at QS promoters. IHF and RNAP-interactions play an important role in LuxR-type regulation in *V. vulnificus* as well (10), suggesting that these mechanisms of gene regulation are conserved across the genus.

Histone-like nucleoid structuring protein (H-NS), which is another nucleoid-organizing protein, functions to directly repress transcription across the genome. H-NS has been best studied in *Escherichia coli* and *Salmonella enterica* (11). At the biophysical level, H-NS is capable of oligomerizing on DNA to form extended filaments and/or DNA-H-NS-DNA bridges (12–14). These nucleoprotein complexes function to impede the activity of RNAP, either by blocking transcription initiation or by inhibiting elongation via topological constraint of the DNA, thereby silencing gene expression from H-NS-bound loci (15, 16). To counter-silence these loci and activate gene expression, bacteria employ transcription factors that are capable of displacing H-NS from promoter DNA. In *V. cholerae*, H-NS modulates the expression of 701 genes (17), and QS-regulated proteins have been postulated to be capable of counter-silencing activities. For example, the QS-controlled protein ToxT is required to activate the *tcpA* and *ctx* promoters, and it is hypothesized that it accomplishes this by displacing H-NS to allow transcription (18).

Here we show that *V. harveyi* LuxR activates transcription of QS genes through anti-repression via H-NS remodeling and/or displacement from QS promoter DNA. RNA-seq and ChIP-seq analyses show that the regulatory overlap between LuxR and H-NS is widespread across the genome. Furthermore, ChIP-qPCR analyses show that H-NS is evicted from QS promoter DNA *in vivo* in a LuxR-dependent fashion. Electrophoretic mobility shift assays coupled with western blots show that LuxR is competent to displace H-NS from promoter DNA *in vitro*. Together, these findings expand on the growing model of LuxR-mediated transcriptional activation to include counter-silencing as a means of activation at H-NS-occupied loci.

## Results

### H-NS is a repressor of bioluminescence in V. harveyi

We previously identified a protein bound to the promoter of the *luxCDABE* bioluminescence genes using a promoter pull-down assay that is predicted to be H-NS (7). The protein encoded by the *hns* gene in *V. harveyi* (*VIBHAR_01827*) shares 53% amino acid identity to *E. coli* H-NS (Fig. S1). Based on the literature that defines H-NS as a transcriptional repressor, we hypothesized that H-NS functions as a repressor of *luxCDABE* at LCD. To test this hypothesis, we constructed a Δ*hns* strain and monitored bioluminescence throughout a growth curve. A typical wild-type bioluminescence growth curve is a “U-shape” (Fig. 1A). At the start of the experiment, stationary HCD cells are diluted into fresh media to LCD. Early in the growth curve, because the cells are at LCD, bioluminescence gene expression is off, due to lack of LuxR, a required activator. The calculation to determine approximate bioluminescence per cell is performed by dividing the bioluminescence by the optical density (OD_600_). Because the bioluminescence number remains constant and low at LCD and the optical density is increasing, the early LCD part of the growth curve has a negative slope. As the population density increases, more LuxR is produced, quorum is reached (approximately at OD_600_ = 0.1), and bioluminescence production increases until stationary phase is reached. A Δ*luxR* strain does not produce bioluminescence and exhibits the negative slope of the data as described throughout the plot (Fig. 1A). Conversely, in a Δ*luxO* strain that produces high levels of LuxR at both LCD and HCD, bioluminescence levels are high throughout the curve (Fig. 1A). The Δ*hns* strain exhibits several important characteristics: 1) the levels of bioluminescence at LCD mirror wild-type, 2) the initial production of bioluminescence appears earlier in growth compared to wild-type, and 3) bioluminescence levels are similar to wild-type at late stationary phase (Fig. 1A). Furthermore, the Δ*luxR* Δ*hns* strain phenocopies a Δ*luxR* strain, indicating that LuxR is epistatic to H-NS. To examine the effect of H-NS copy number on bioluminescence, we developed a high-throughput bioluminescence assay using a plate reader that monitors bioluminescence during the latter part of the “U-shape” curve. We complemented the *hns* deletion using an ectopic IPTG-inducible expression vector and observed bioluminescence levels comparable to wild-type (Fig. 1B). Using an array of IPTG concentrations, we observed dose-responsiveness between H-NS expression and bioluminescence; at 50 µM IPTG, bioluminescence is severely decreased (Fig 1B). To determine whether *V. harveyi* H-NS functions similarly to *E. coli* H-NS, we complemented the Δ*hns V. harveyi* strain with the *hns* gene from *E. coli* (*hns_Ec_*). Expression of *hns_Ec_* complemented the bioluminescence phenotype in the Δ*hns* strain to produce approximately wild-type levels of bioluminescence (Fig. 1C). At 50 µM IPTG, expression of *E. coli* H-NS results in bioluminescence levels that are less than wild-type. In addition, the L30P allele of *E. coli* H-NS is able to bind DNA but unable to oligomerize, which is essential for its repressor functionality *in vivo* (19, 20). We therefore complemented the Δ*hns* strain with *V. harveyi hns* L30P allele to test its activity. As expected, the Δ*hns* strain expressing H-NS^L30P^ phenocopied the Δ*hns* strain and produced more bioluminescence than wild-type during exponential phase (Fig. 1D). Even at high concentrations of IPTG, this mutant is unable to complement the Δ*hns* strain. From these data, we conclude that *V. harveyi* H-NS represses bioluminescence in a concentration dependent manner and is a functional homologue of H-NS from *E. coli*. Further, the role of H-NS in repression of *luxCDABE* occurs at HCD; at LCD, deletion of *hns* has no impact on bioluminescence.

**Figure 1.**
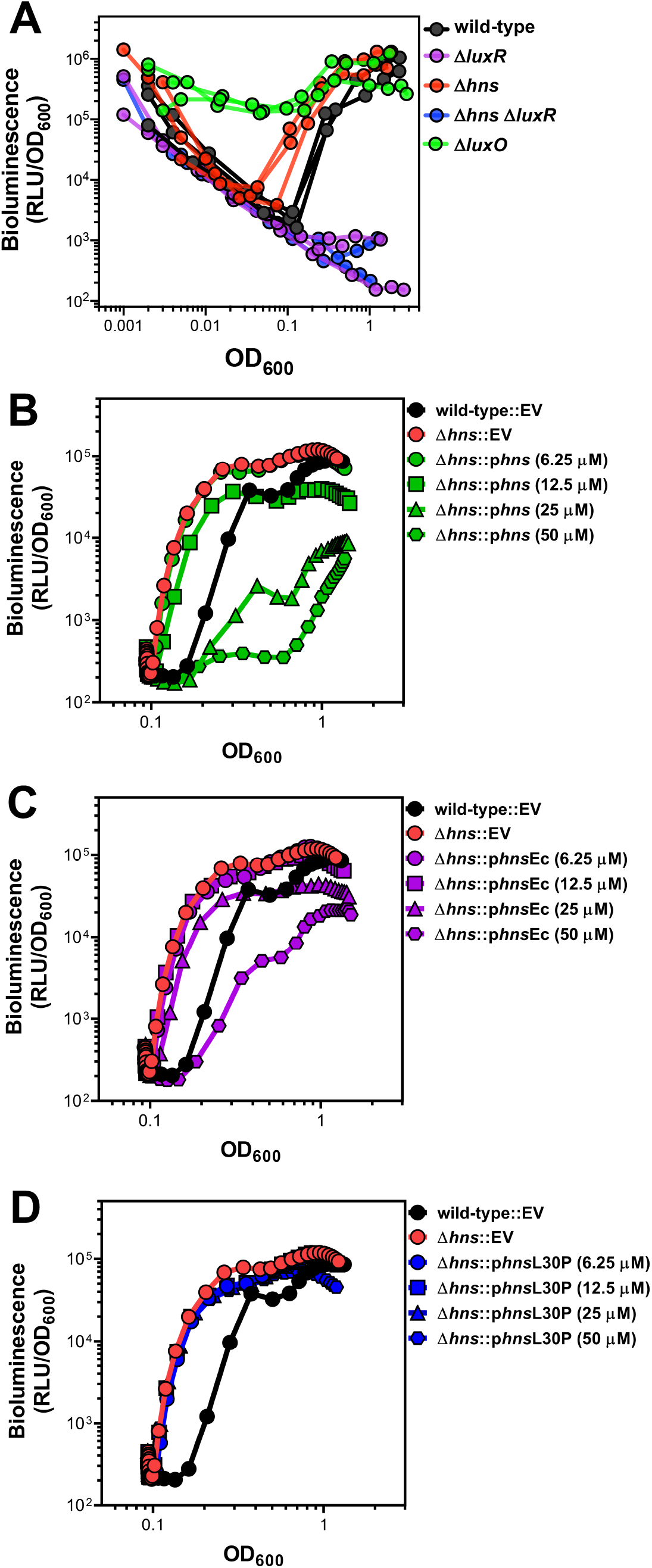
H-NS controls bioluminescence in *V. harveyi* and is functionally similar to H-NS in *E. coli*. (A) Bioluminescence (relative light units, RLU; Lux/OD_600_) was measured for *V. harveyi* wild-type (BB120), Δ*luxR* (KM669), Δ*hns* (RRC045), Δ*hns* Δ*luxR* (RRC168), and Δ*luxO* (JAF78). Data shown are three independent biological experiments. (B, C, D) Bioluminescence (relative light units, RLU; Lux/OD_600_) was measured for *V. harveyi* wild-type (BB120) and Δ*hns* (RRC045) strains. Strains contained either an empty vector (EV, pMMB67EH-kanR), a plasmid expressing *V. harveyi hns* under an IPTG-inducible promoter (pRC053; panel B), a plasmid expressing *E. coli hns* under an IPTG-inducible promoter (pRC054; panel C), or a plasmid expressing *V. harveyi hns* L30P under an IPTG-inducible promoter (pRC055; panel D). IPTG was added at concentrations of 6.25 μM, 12.5 μM, 25 μM, or 50 μM; IPTG concentration had no effect on the curve for the wild-type EV or Δ*hns* EV strains (data not shown). Data shown are averages of two technical replicates and are representative of five independent biological experiments.

### H-NS and LuxR co-regulate 124 genes

To determine whether other promoters are regulated by H-NS and LuxR in a similar manner to P_*luxCDABE*_, we used RNA-seq to define the regulons for LuxR and H-NS. For the RNA-seq, we compared transcripts in a Δ*hns* strain to a wild-type strain. Conventionally, we have performed QS-related RNA-seq analyses at OD_600_ = 1.0 because this is when HCD is reached. Therefore, we also performed the wild-type vs. Δ*hns* RNA-seq at this point in the growth curve. We observed that H-NS regulates 726 genes (>2-fold, *p* < 0.05; Fig. 2A, dataset S1) in *V. harveyi*, which is strikingly comparable to the 701 genes regulated by H-NS in *V. cholerae* (21). Among the 726 genes, 596 are repressed by H-NS. Because H-NS is a global repressor and thus functions in the opposing manner to LuxR at *luxCDABE*, we hypothesized that LuxR acts to derepress H-NS-bound loci. Indeed, the *luxCDABE* genes were absent from the H-NS regulon in the wild-type background, which supports this hypothesis. To determine if the presence of LuxR affects H-NS gene regulation at other loci, we next performed RNA-seq comparing a Δ*luxR* strain to a Δ*luxR* Δ*hns* strain. Under these conditions, H-NS regulates 824 genes (>2-fold, *p* < 0.05; Fig. 2A, dataset S1) and, as expected, the *luxCDABE* genes were present. However, only 430 genes are repressed by H-NS in this comparison. Together, these RNA-seq data show that H-NS regulates a similar number of genes in the presence or absence of LuxR, though the scale and direction of regulation varies widely. Among the combined genes in the H-NS regulons (wild-type and Δ*luxR* backgrounds combined), 178 genes overlapped with the previously identified 308-gene LuxR regulon (>2-fold, *p* < 0.05; Fig. 2A; (7)). These genes include many systems previously shown to be highly regulated by LuxR, including the *luxCDABE* genes, the type III secretion operons, the *proXWV* glycine betaine transport system, and several ABC transport systems, among others. From these data, we conclude that H-NS and LuxR are relevant global co-regulators in *V. harveyi*.

**Figure 2.**
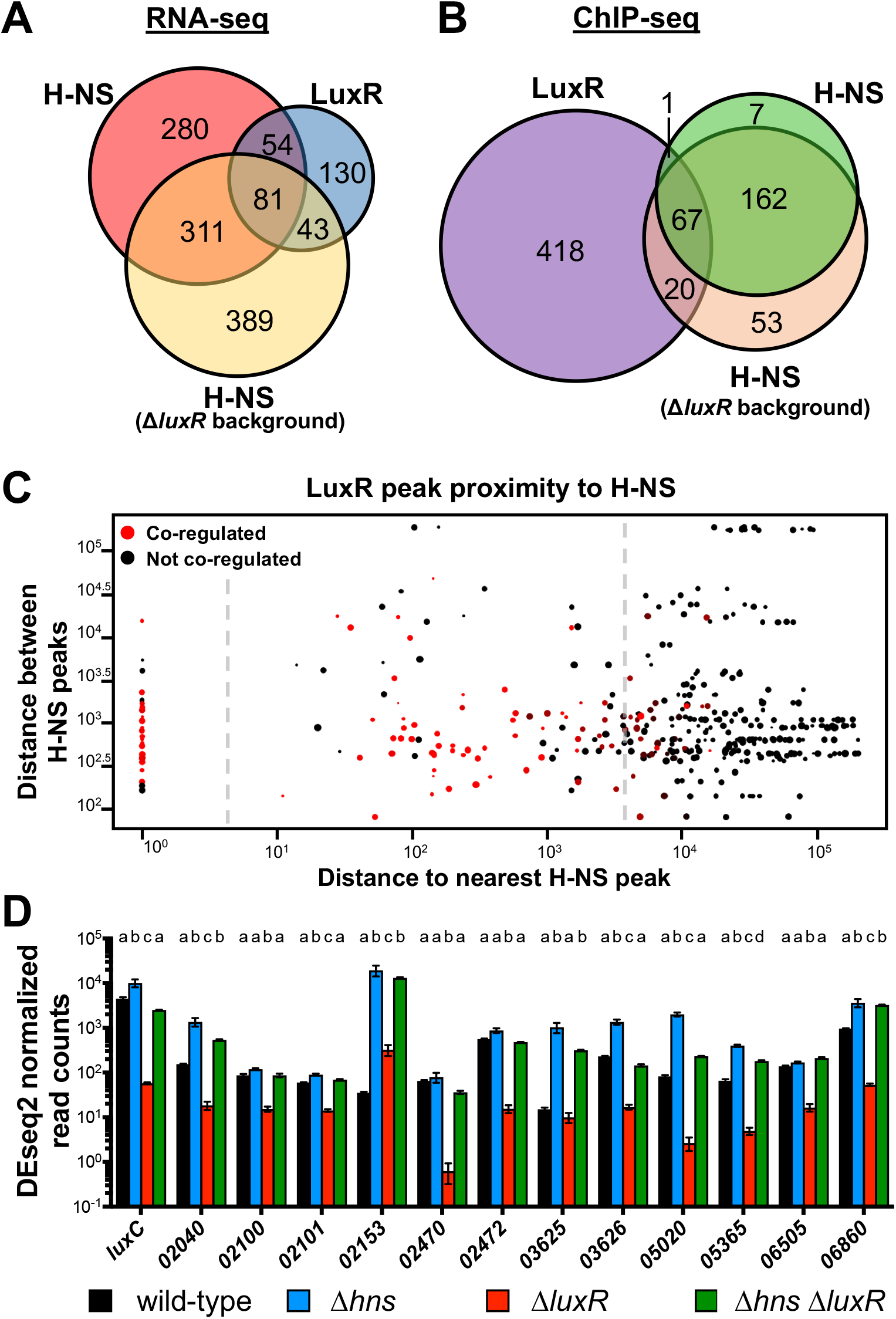
LuxR and H-NS directly co-regulate QS genes. (A) RNA-seq was used to determine genes regulated >2-fold, *p <* 0.05. The H-NS regulon was determined by comparing transcript levels of a Δ*luxR* strain (KM669) to a Δ*hns* Δ*luxR* strain (RRC168). The LuxR regulon was previously determined by comparing wild-type (BB120) to a Δ*luxR* strain (KM669) (7). (B) FLAG-LuxR and H-NS-FLAG ChIP-seq profiles were analyzed for overlapping peaks within 500bp. The H-NS-FLAG ChIP-seq profile was determined in the Δ*luxR* background (RRC237), and the FLAG-LuxR ChIP-seq profile was determined in a wild-type background (JV039). (C) Each point represents a LuxR binding site with the diameter corresponding to the strength of the peak at that site; all 506 LuxR peaks are plotted. The distance between a LuxR peak and the nearest H-NS peak is plotted on the x-axis. The distance between the nearest H-NS peak and the next proximal H-NS peak is plotted on the y-axis. Red points represent LuxR peaks that are proximal to genes that are co-regulated by LuxR and H-NS while black points represent genes that are regulated by only one protein or neither. (D) DESeq2 normalized read counts determined by RNA-seq of RNA collected from wild-type (BB120), Δ*luxR* (KM669), Δ*hns* (RRC045), Δ*hns* Δ*luxR* (RRC168) *V. harveyi* strains. *V. harveyi* locus tags are listed on the x-axis (*VIBHAR_XXXXX*). Different letters indicate significant differences between strains (*p <* 0.05; two-way analysis of variation (ANOVA), followed by Tukey’s multiple comparison test of log-transformed data; *n=3*).

### LuxR and H-NS co-occupy and co-regulate 19 promoters

Based on our bioluminescence and transcriptomic data, our working hypothesis was that as LuxR protein concentration increases during QS, LuxR binds to promoters containing LuxR binding sites to displace H-NS, which alleviates promoters from H-NS-mediated repression. To test this hypothesis and identify promoters that are bound by both LuxR and H-NS, we used ChIP-seq to define global binding profiles for LuxR and H-NS individually. *FLAG*-*luxR* and *hns-FLAG* unmarked alleles were each introduced separately into *V. harveyi* BB120, and ChIP-seq was performed comparing each of the FLAG-tagged mutant strains to the wild-type strain. We identified 506 binding peaks for LuxR and 237 binding peaks for H-NS across the genome (Fig. 2B, dataset S2). We also analyzed LuxR and H-NS binding peaks to determine how many are proximal to each other and found that LuxR and H-NS peaks overlap at 68 regions (Fig. 2B, dataset S2). Furthermore, we analyzed the distance between H-NS and LuxR peaks across the genome. We observed distinct types of LuxR binding site locations using this analysis: 1) LuxR binding sites that directly overlap with H-NS binding sites, 2) LuxR binding sites that are proximal to H-NS binding sites (within ∼1,000 bp), and 3) LuxR binding sites that are distant from H-NS peaks (>5,000 bp away) (Fig. 2C). Interestingly, most LuxR peaks within 1,000 bp of an H-NS peak are proximal to genes that are co-regulated by both LuxR and H-NS (red dots, Fig. 2C). As expected, we observed LuxR and H-NS peaks at the *luxCDABE* promoter (Fig. 3A), and no peaks at a control locus (*clpP*; Fig. 3B).

**Figure 3.**
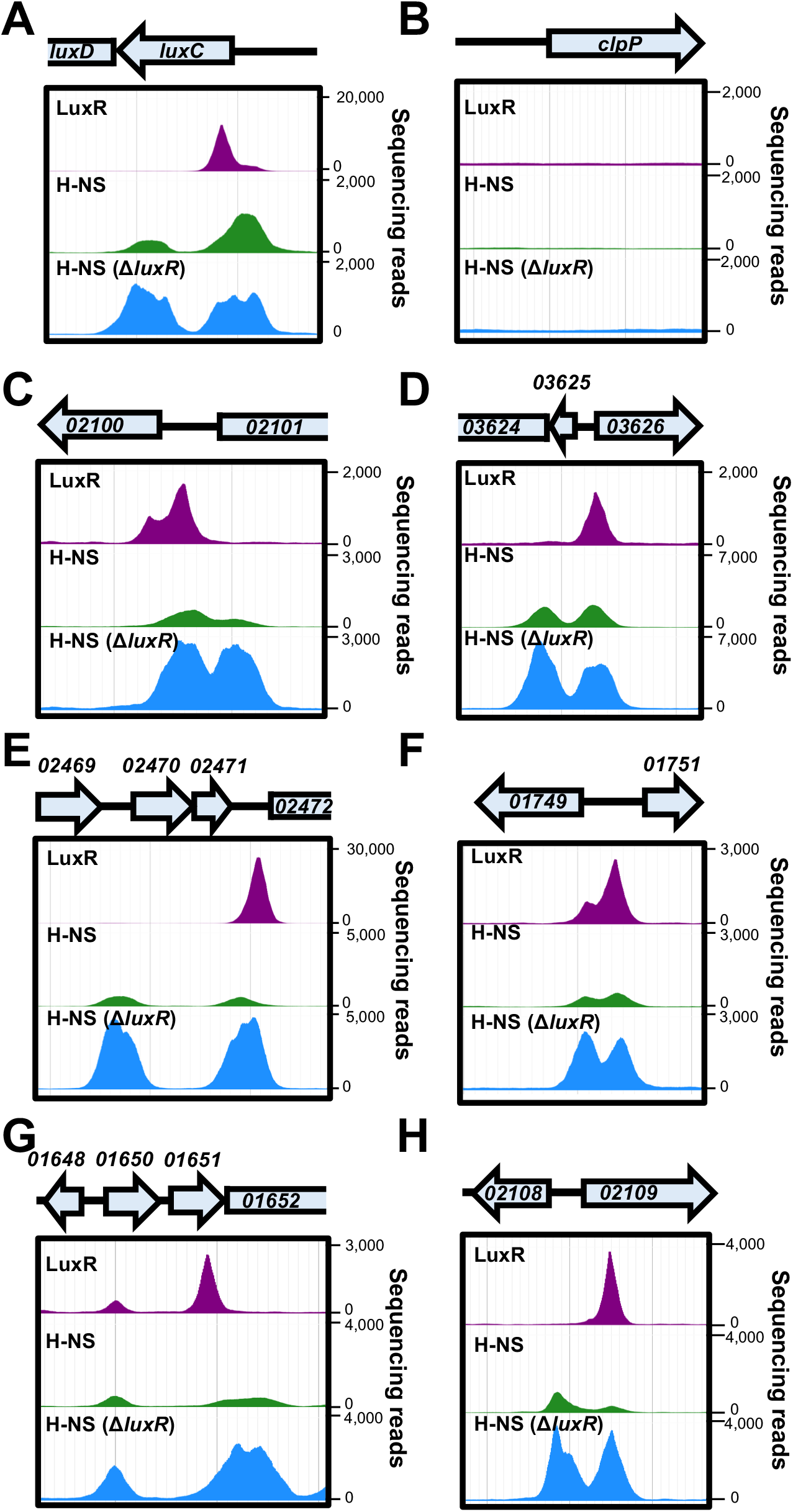
LuxR and H-NS binding peak distribution. (A-H) LuxR and H-NS ChIP-seq peaks at QS loci *luxC*, *VIBHAR_02100*, 03626, 02472, 01749, 01652, and 02108 are shown for experiments in which LuxR (JV039) or H-NS (RRC169) were ChIPed in the wild-type background (top two rows for each panel) or in the Δ*luxR* background (RRC237, bottom row for each panel). ChIP-seq profiles were visualized using JBrowse 1.12.3.

The ChIP-seq experiments described above were performed with cells collected at HCD (OD_600_ = 1.0), similar to our RNA-seq analyses because this is the time point is when LuxR protein is maximally produced prior to stationary phase (5). Based on our hypothesis that LuxR displaces H-NS at HCD, we reasoned that our experimental ChIP-seq setup may have missed H-NS binding sites that were obscured by competition with LuxR because they were collected at the HCD state. Therefore, to observe all H-NS-bound sites in the genome, we performed ChIP-seq with H-NS-FLAG in a Δ*luxR* mutant strain background. As we predicted, the number of H-NS binding peaks increased from 237 to 302 in the absence of LuxR (dataset S2). The number of co-occupied H-NS and LuxR loci increased from 68 to 87 in the absence of LuxR (dataset S2). By examining these 87 regions further in the context of transcriptomic data and gene organization, we narrowed this list down to 65 regions that are co-occupied by LuxR and H-NS (dataset S2). Among these are 28 promoters controlling expression of 63 genes that are also co-regulated by LuxR and H-NS (dataset S2). Examples of binding peaks at co-regulated and/or co-occupied regions are shown in Figure 3. The remaining co-occupied loci are not present in the LuxR/H-NS co-regulon. These genes may be regulated <2-fold, regulated by only one of these proteins, or regulated only under specific conditions that we did not test. From these data, we conclude that H-NS and LuxR co-localize throughout the genome to co-regulate genes.

### LuxR binding alters H-NS binding profiles *in vivo*

From the ChIP-seq data, we also observed substantial differences in H-NS peak heights in the presence or absence of LuxR. We hypothesized that these profile changes are due to binding of LuxR and displacement of H-NS. To ensure peak height differences were not due to differences in sample preparation (*i.e.* library preparation/number of sequencing reads), we identified 98 loci that contain H-NS peaks but are devoid of any LuxR peaks. We reasoned that these peaks would not be directly influenced by LuxR and thus we could normalize the H-NS sequencing read distribution to these peaks (normalization procedure detailed in methods). Following normalization, we re-compared these 98 peaks and found that the loci remained significantly correlated between wild-type and Δ*luxR* backgrounds (Fig. S5). With the read counts normalized, we then analyzed the H-NS peak heights at LuxR-bound loci. Each of the loci which are directly activated by LuxR/repressed H-NS showed an increase in H-NS peak height in the absence of LuxR (Fig. 3A, 3C, 3D, 3E, 3F), indicating increased binding of H-NS at these loci. Interestingly, when we analyzed the difference in H-NS peak heights between the wild-type and Δ*luxR* backgrounds, we noticed a single LuxR binding site often influenced both H-NS peaks, even if separated by 2 kb as is the case with *VIBHAR_02472* (Fig. 3E). Even at two loci that LuxR does not regulate under our tested conditions, there is a measurable H-NS peak height difference when LuxR is bound compared to the Δ*luxR* background (Fig. 3G, 3H).

Another intriguing property of the H-NS ChIP-seq profile is the frequent occurrence of doublets in which two peaks of similar height are proximal to one another. This phenomenon is also detectable in H-NS ChIP-seq experiments performed in *V. cholerae* (22). We suspect that these peak profiles represent bridges formed by H-NS. While the peak height is generally similar within doublets, the distance between peaks is highly variable ranging from ∼400 bp to 2 kb. We analyzed the H-NS ChIP-seq data for binding sites that matched these parameters (2 similarly sized peaks within 2 kb of each other). We identified 52 H-NS doublets throughout the genome, and 32 are proximal to a LuxR binding site (within 1 kb). The notable disruption of these H-NS doublets by the presence of LuxR (*e.g.*, Fig. 3A, 3C, 3H) suggests that LuxR influences H-NS binding distribution, and thus H-NS bridging. Collectively, these data suggest that LuxR binding displaces H-NS, decreasing H-NS binding and/or bridging at co-regulated and co-bound loci.

### LuxR is not required for transcription in the absence of H-NS

Based on our observations of LuxR activation and H-NS repression activities at the *luxCDABE* promoter, our working hypothesis was that as LuxR protein concentration increases during QS, LuxR binds to P_*luxCDABE*_ and displaces H-NS, which alleviates promoters from H-NS-mediated repression. As a control, we examined gene expression of the *luxCDABE* genes to compare to our bioluminescence assays and previously published expression data. The RNA-seq data for the *luxC* gene indicate that Δ*hns* cells express ∼2-fold more *luxC* transcripts than wild-type cells at OD_600_ = 1.0 (Fig. 2D). As expected, the Δ*luxR* strain exhibits very low levels of *luxC* transcript (Fig. 2D), which has been shown previously (23) and is evidenced by lack of bioluminescence production (Fig. 1A). We were therefore surprised to observe that the Δ*luxR* Δ*hns* strain restored *luxC* transcripts with only ∼2-fold fewer transcripts compared to wild-type and ∼4-fold fewer than the Δ*hns* strain (Fig. 2D). This result suggests that LuxR is only required for *luxC* transcription in the presence of H-NS. However, the bioluminescence production of the Δ*luxR* Δ*hns* strain remains very low (Fig. 2D), suggesting that a posttranscriptional mechanism of regulation coordinated via LuxR is required for bioluminescence production.

To test whether this occurs at other LuxR/H-NS co-regulated and co-occupied promoters, we examined the other 9 loci in addition to *luxCDABE* that are activated by LuxR and repressed by H-NS (dataset S2). Each of the 37 genes under control of the 13 promoters show the same pattern of gene expression as *luxC*: transcript levels are lowest in the Δ*luxR* strain for all genes, suggesting that LuxR is required for activation of these genes (Fig. 2D). However, when *hns* is deleted in the Δ*luxR* background, transcript levels are completely or partially restored to wild-type levels (Fig. 2D). From these data, we conclude that LuxR is not required for transcription activation of these promoters in the absence of H-NS.

### H-NS is evicted from promoter regions as cells transition from LCD to HCD

To investigate the role of LuxR in H-NS localization at promoters, we developed a ChIP-qPCR assay that mimics various QS states to quantify relative amounts of LuxR and H-NS protein occupancy at promoters. These experiments were performed using a strain background in which the genes encoding the HAI-1 autoinducer synthase and CAI-1 and AI-2 receptors were deleted (TL25: Δ*luxPQ*, Δ*luxM*, Δ*cqsS*) (5). This strain responds solely to exogenously supplied HAI-1. Based on previously established concentrations of HAI-1 that mimic the transition states between LCD and HCD, we tested a range of synthetic HAI-1 from 0-1000 nM. When either no HAI-1 or low concentrations of HAI-1 are added to this strain, both *aphA/luxR* transcripts and AphA/LuxR proteins are detected (5). As the HAI-1 concentration added increases to 1000 nM, *aphA* transcript sharply decreases and AphA protein is no longer detected, whereas *luxR* transcript and LuxR protein increase (5). To pinpoint AI concentrations that mimic LCD, mid cell density (MCD), and HCD, a bioluminescence induction assay was used. Based on the bioluminescent output, we found that 0/4 nM, 40 nM, and 1000 nM were sufficient to replicate light production levels present at LCD, MCD, and HCD (Fig. S2). The *FLAG-LuxR* and *H-NS-FLAG* alleles were individually introduced into the TL25 background, and ChIP was performed using these strains grown with the above HAI-1 concentrations. The immunoprecipitated DNA was analyzed via qPCR directed at 7 promoters that showed strong, overlapping peak signals for LuxR and H-NS. The promoter region upstream of *clpP* was used a negative control for this experiment because no ChIP-seq peaks were detected for LuxR/H-NS (Fig. 3B), and similarly, this promoter is not captured by LuxR or H-NS with the ChIP-qPCR assay (Fig. 4B). At promoters in which LuxR and H-NS are both present, such as P_*luxC*_, LuxR protein occupancy increases at the promoter while H-NS occupancy decreases as cells transition from LCD to HCD (Fig. 4A). This trend was conserved at 6 additional promoters in which we observed a marked increase in LuxR occupancy with a simultaneous decrease in H-NS occupancy (Fig. 4C-H). While the overall trend of decreasing H-NS occupancy is consistent across promoters, the amount of HAI-1 required to reach maximum H-NS displacement is variable. Some promoters, such as P_*luxC*_ and P_02100_, require 1000 nM HAI-1, whereas others, such as P_01749_ and P_02472_, require ≤40 nM HAI-1. Taken together, these data support the hypothesis that LuxR accumulation coincides with the displacement of H-NS from QS promoters.

**Figure 4.**
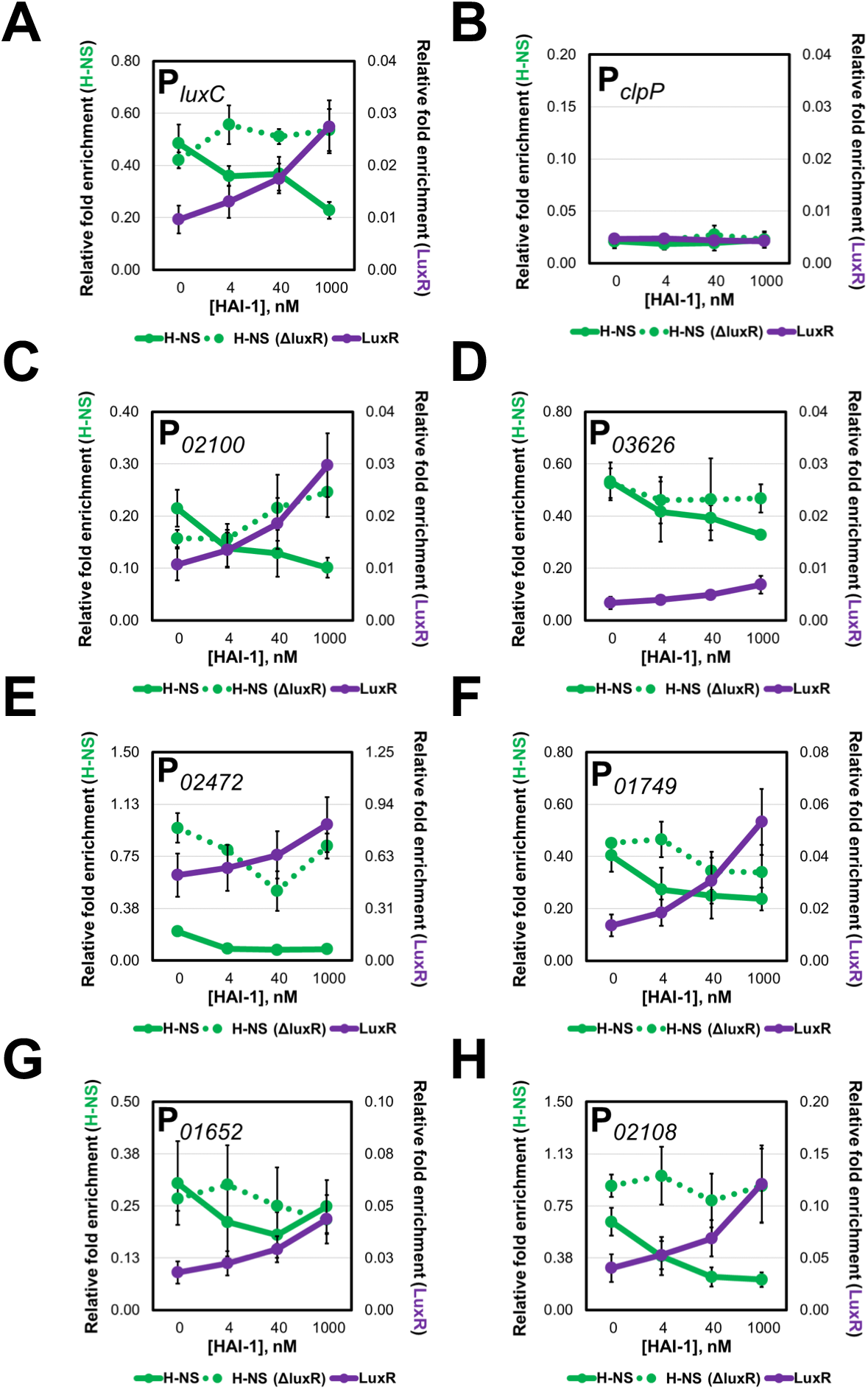
LuxR and H-NS occupancy is inversely related at QS loci. (A-H) The relative enrichment of DNA from various promoters analyzed by ChIP-qPCR experiments in the presence of 0, 4 nM, 40 nM, or 1000 nM HAI-1 in *hns-FLAG* (RRC169), *FLAG-luxR* (JV039), and *hns-FLAG* Δ*luxR* (RRC237) strains.

To test if LuxR is necessary and/or sufficient to displace H-NS from QS loci *in vivo*, we constructed the TL25 FLAG-*hns* Δ*luxR* strain. We performed our ChIP-qPCR analysis to monitor changes in H-NS occupancy in the absence of LuxR. We observed that the relative amount of H-NS present at a promoter is higher in the Δ*luxR* background compared to wild-type at some or all concentrations of HAI-1 (Fig. 4B-H). While overall H-NS occupancy is higher in these profiles, we do observe H-NS displacement in the absence of LuxR at certain promoters. For instance, at P_02472_ H-NS displacement is observed between 0-40 nM HAI-1 and at P_01749_ H-NS is displaced between 4-40 nM HAI-1. These results are not surprising as LuxR likely works alongside other transcription factors to activate transcription; thus, the observation of H-NS displacement in the absence of LuxR can be attributed to the action of other regulators. The results of these experiments provide evidence that LuxR is required for maximal H-NS displacement at QS promoters.

### LuxR displaces H-NS from promoter DNA in vitro

The results of the ChIP-qPCR experiments showed that increasing concentrations of AI and LuxR-binding results in lower H-NS binding at QS promoters *in vivo*. To determine whether LuxR is able to displace H-NS from DNA *in vitro*, we first used a bioinformatics approach (FIMO software, (24)) to identify putative H-NS binding sites in the *luxCDABE* promoter. Using the high-affinity H-NS nucleation site identified in *E. coli* (TCGATAAATT) (25), we identified 13 potential sites in P_*luxCDABE*_, many of which overlap with the seven established LuxR binding sites (Fig. 5A, Table S1). Next, we used electrophoretic mobility gel shift assays (EMSAs) and performed titration curves with purified 6xHis-H-NS (referred to as H-NS) and LuxR individually to determine the binding kinetics for each protein with *luxCDABE* promoter DNA. For both sets of reactions, we observed the DNA shift to a larger molecular weight, suggesting that all the DNA was bound (Fig. 5B). For H-NS, a steep binding curve was observed, which likely reflects the cooperative DNA binding activities of H-NS (26, 27). For LuxR, multiple intermediate LuxR-DNA bands were observed, as expected (7).

**Figure 5.**
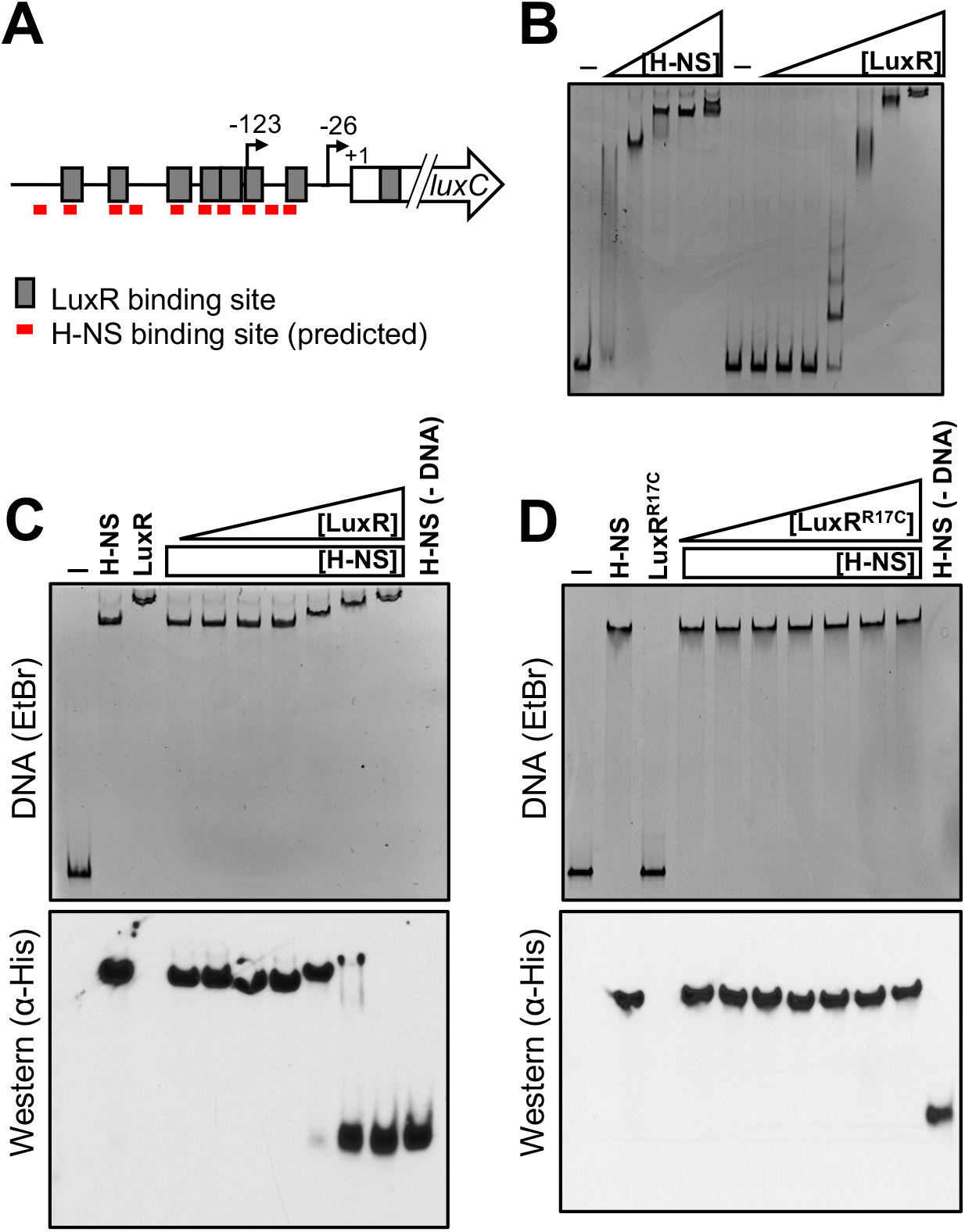
H-NS is displaced from P_*luxC*_ DNA by LuxR *in vitro*. (A) Diagram of *luxCDABE* promoter. Gray boxes indicate LuxR binding sites, and red lines indicate FIMO-predicted H-NS binding sites. Arrows indicate transcription start sites relative to the start codon (+1). (B) EMSA reactions containing 8.8 nM P_*luxC*_ DNA and either H-NS (100, 150, 200, 250, 300 nM) or LuxR (31.25, 62.5, 125, 250, 500, 1000, 2000 nM) purified protein. Reactions denoted ‘-’ contained no protein. (C) Competitive EMSA reactions consisting of 8.8 nM P_*luxC*_ DNA and 200 nM purified H-NS and either 31.25, 62.5, 125, 250, 500, 1000, or 2000 nM purified LuxR. Lanes labelled ‘H-NS’, ‘LuxR’, ‘-’, and H-NS (−DNA) contained 200 nM H-NS, 2000 nM LuxR, no protein, and 200 nM H-NS without DNA, respectively. The reactions were run on polyacrylamide gels and stained with ethidium bromide (top) and then transferred to a nitrocellulose membrane and probed for H-NS using α-His-HRP antibodies (bottom). (D) The same experimental procedures were used as described in panel C but instead using the DNA-binding deficient LuxR^R17C^ purified protein.

To examine competition between LuxR and H-NS, we generated reactions with both H-NS and LuxR added to *luxCDABE* promoter DNA. We added a constant amount of H-NS and varying concentrations of LuxR. We observed that increasing concentrations of LuxR incrementally shifted the migration of the nucleoprotein complex to mirror that of the LuxR-P_*luxCDABE*_ complex (Fig. 5C). This result suggests that LuxR evicts H-NS from this promoter. To determine whether any H-NS protein remained bound to the shifted DNA substrate, EMSA reactions were then transferred to a nitrocellulose membrane such that the proteins associated with bound DNA could be monitored via western blot. We observed that as LuxR concentration increases across reactions, the H-NS protein signal corresponding to the H-NS-P_*luxCDABE*_ bound complex decreases. In parallel, the signal corresponding to unbound H-NS accumulates (Fig. 5C).

The experiment described above was performed in the absence of competitor DNA because H-NS can bind to DNA non-specifically. To verify that the displacement of H-NS was dependent on LuxR associating with specific binding sites rather than non-specific interactions with the DNA, we performed the same competition experiment in the presence of competitor DNA (poly[dI-dC]) to limit the amount non-specific protein-DNA interactions. Under these conditions, LuxR remains able to displace H-NS from P_*luxCDABE*_ DNA (Fig. S3C). Importantly, we tested a LuxR^R17C^ DNA binding mutant (8) and found that it was unable to displace H-NS from the *luxCDABE* promoter (Fig. 5D). This result confirms that LuxR eviction of H-NS relies on LuxR DNA binding activity. Next, we extended these experiments to two additional promoters that are repressed by H-NS and activated by LuxR (P_02100_ and P_03626_). Competition between H-NS and LuxR was observed using these promoters as well (Fig. S3A, S3B), indicating that LuxR eviction of H-NS is a conserved mechanism at multiple loci.

Lastly, we wanted to examine the influence of H-NS on the binding of RNAP to P_*luxCDABE*_ promoter DNA. Using the same competitive EMSA experimental design as described above, we pre-bound the P_*luxCDABE*_ substrate with H-NS and titrated in various amounts of purified *E. coli* 10xHis-RNAP (referred to as RNAP). With increasing concentrations of RNAP, the H-NS-DNA complex showed slower migration through the gel (Fig. S4). This observation indicates that RNAP is capable of binding H-NS-bound P_*luxCDABE*_ promoter DNA. However, although RNAP can bind the H-NS-bound DNA, it is not sufficient to displace detectable amounts of H-NS (Fig. S4). Together, these results indicate two important properties of H-NS: 1) LuxR is capable of displacing H-NS through competitive DNA binding, and 2) RNAP can bind to H-NS-bound DNA but cannot displace H-NS at the *luxCDABE* promoter.

## Discussion

LuxR/HapR have been extensively studied for decades as key regulators of QS genes in vibrios, yet our collective understanding of the molecular workings of these unique transcriptional regulators has remained limited. In the present study, we show that LuxR engages in a functional relationship with H-NS in addition to IHF and RNA polymerase. We show that the actions of LuxR and H-NS are fundamentally opposed in order to coordinate LCD/HCD gene expression programs in *V. harveyi*.

Derepression describes a transcriptional regulatory mechanism in which a negative regulator is cleared from a promoter to enable the recruitment and/or stabilization of RNAP and thereby stimulate transcription. H-NS has been the subject of studies investigating derepression mechanisms in Gram-negative bacteria. Similarly to *V. cholerae* ToxT, the positive regulator LeuO evicts H-NS from the DNA at the *ompS1* promoter in *Salmonella enterica* (28). The work presented in this study demonstrates that LuxR is also capable of derepression of H-NS to directly activate gene expression. At one example promoter, the *luxCDABE* bioluminescence promoter, our experiments provide compelling evidence for a model in which H-NS oligomerizes to form a filament and/or bridge that represses gene expression. The *V. harveyi* H-NS L30P mutant allele fails to complement bioluminescence in the Δ*hns* strain in *V. harveyi*, which strongly suggests that oligomerization (*i.e.* filament formation) is essential for H-NS to serve as a negative regulator at the *luxCDABE* promoter. Because the DNA-binding deficient LuxR^R17C^ mutant was unable to displace H-NS *in vitro*, this suggests that LuxR evicts H-NS from DNA through competition for the same binding sites. Notably, because both LuxR and H-NS DNA binding sites are AT-rich sequences, it is unsurprising that they may compete for similar sites at a promoter.

Within the 506 and 302 peaks identified for LuxR and H-NS, respectively, 87 peaks overlapped and were proximal to an ORF (≤500 bp). The LuxR ChIP-seq profile previously published identified 1165 LuxR binding peaks (8). However, that previous experiment was performed with *FLAG*-*luxR* expressed from a multi-copy plasmid in at Δ*luxR* background, which is the likely reason for the increased number of LuxR binding peaks identified. Overall, the ChIP-seq data show that QS loci are regulated via the competitive binding dynamics of LuxR and H-NS (and likely other regulators). The function of LuxR at 13 of these promoters appears to be eviction of H-NS, and this results in activation of gene expression. The nucleoprotein bridges formed by H-NS may inhibit transcription initiation, trap elongating RNAP, and/or affect the topology of the DNA such that RNAP elongation is inhibited. Importantly, we observed that at many promoters, the two H-NS peaks were positioned such that the promoter was between the peaks. If these peaks indeed represent DNA-H-NS-DNA bridges, it is likely that even if RNAP can initiate transcription effectively, the bridge would block elongation. In addition, our finding that several H-NS peaks colocalize near genes repressed by LuxR presents the question: does LuxR also function alongside H-NS to repress gene expression? Additional experimentation will be necessary to properly address this question, but we speculate that LuxR may be sufficient to displace transcriptional activators that prevent H-NS from binding to these loci. In support of this theory is the finding that HapR and H-NS function together to ‘double-lock’ the *vieSAB* promoter in *V. cholerae* (29).

Perhaps the most surprising evidence that LuxR functions to oppose the action of H-NS globally is the observation that transcript levels of many QS genes are restored to wild-type levels in the Δ*luxR* Δ*hns* strain. This finding implies that, at HCD, LuxR becomes dispensable when H-NS is not present for a subset of genes. It is important to note that, for at least *luxCDABE*, while transcript levels are restored to near wild-type levels in the Δ*luxR* Δ*hns* strain, the bioluminescence phenotypic output remains very low. This result suggests that the bioluminescence genes are controlled both transcriptionally and post-transcriptionally. One possibility is that LuxR regulates the expression of a sRNA or an RNA-binding protein that is necessary for translation of these genes. Alternatively, a *cis*-acting RNA regulatory element such as a riboswitch or RNA thermometer may be regulating the translation of the *luxCDABE* mRNA. Further analyses will be required to test these possibilities.

From the collective work on the *luxCDABE* promoter, it is clear that these genes are highly regulated by multiple mechanisms, likely due to the cost of expressing these proteins at such high levels. At LCD, the ChIP-seq H-NS peak profile suggests that H-NS is bound to the *luxCDABE* promoter in a nucleoprotein bridge. LuxR protein levels are insufficient to stimulate transcription of the *luxCDABE* genes, even in the absence of *hns*. Thus, there are likely other regulatory mechanisms acting at the transcriptional and/or posttranscriptonal levels to repress expression at LCD. As population density increases and LuxR protein levels rise, LuxR binds to the binding sites in the *luxCDABE* promoter, and the H-NS nucleoprotein filament is destabilized resulting in the mass eviction of H-NS monomers/dimers from the DNA. Once the promoter has been purged of H-NS, it is probable that LuxR and other transcription factors, such as IHF, remodel promoter architecture to poise RNAP for transcription of the *luxCDABE* genes. LuxR interaction with RNAP has been established at the *luxCDABE* promoter (9), and likely is responsible for the ∼4-fold increase in transcription in the presence of LuxR. The observation that RNAP can bind to H-NS-bound DNA but is unable to displace H-NS is suggestive that RNAP recruitment is not the rate-limiting step for expression of the *luxCDABE* operon. Instead, it is likely that RNAP can bind to the *luxCDABE* promoter but remains incompetent for transcription. This hypothesis aligns well with the observation that RNAP can partially elongate through H-NS-DNA bridges but inevitably pauses due to the topological constraints of the H-NS-DNA bridge (13). Additionally, H-NS filaments can interact with promoter-bound RNAP to inhibit open complex formation (30). While the experiments performed in this study are insufficient to identify which mechanism(s) occurs at LuxR-dependent promoters, we speculate that the mechanism is probably promoter dependent.

We therefore propose an updated model for the activation of QS genes (such as the *luxCDABE* operon) in *V. harveyi* (Fig. 6). At LCD, the amount of LuxR in the cell is inadequate to compete against H-NS for binding of QS promoter DNA. Under these conditions, H-NS remains bound to QS promoters as a nucleoprotein bridge, thereby silencing gene expression. As the population transitions to HCD, the intracellular concentration of LuxR increases ∼10-fold (31). Given the gradual accumulation of LuxR as well as the variation of binding site affinities, LuxR likely associates with high-affinity binding sites followed by moderate to low affinity sites within promoter regions. We propose that this incremental binding of LuxR to the promoter is part of a stepwise mechanism that allows for the systematic displacement of H-NS nucleoprotein bridges and filaments. Once H-NS bridges have been remodeled and H-NS displaced from the promoter, LuxR aids in the recruitment and/or stabilization of RNA polymerase to increase transcription levels (9). Thus, in a Δ*hns* strain, LuxR is not required for transcription, but the presence of LuxR increases transcription by interaction with RNA polymerase.

**Figure 6.**
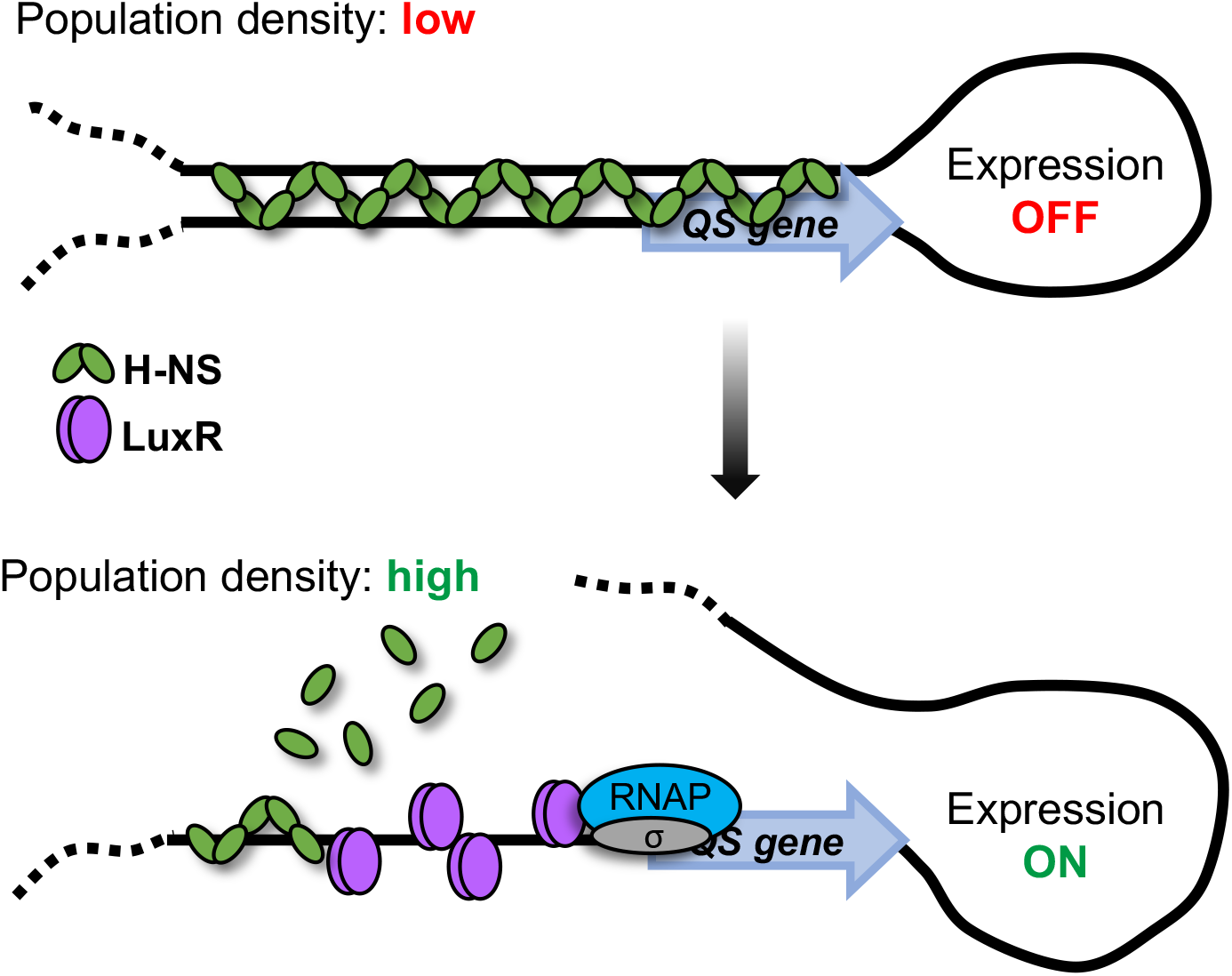
QS promoters are activated through LuxR-dependent displacement of H-NS. At low cell density, the levels of LuxR are low and thus H-NS outcompetes LuxR for binding DNA at QS promoters. In this state, the DNA-H-NS-DNA bridge blocks transcription and the gene is not expressed. As cells transition to high cell density, LuxR concentration in the cell increases, which enables LuxR to outcompete H-NS for binding of QS promoter DNA and disrupt inhibitory H-NS bridges and filaments. These events together with LuxR interaction with RNA polymerase alpha drives maximal transcription of QS genes.

Collectively, biochemical and genetic studies of LuxR/HapR proteins have made significant progress in determining the mechanism of transcriptional activation and repression by these master QS regulators. LuxR-type proteins function alongside IHF, directly oppose H-NS, associate with RNAP, and stimulate and/or block activities of other transcription factors (4,7,9,10,32–34). Importantly, though many of these mechanisms are conserved in *Vibrio* species and at multiple promoters, we surmise that the transcriptional control of each QS promoter is likely different. The *luxCDABE* promoter provides an easily monitored system for examining possible LuxR functions that can be tested at other promoters. Our observation that LuxR counter-silencing of H-NS is globally functional in *V. harveyi* and that similar observations have been reported in other QS bacteria alludes that H-NS counter-silencing via QS regulators could represent a conserved QS regulatory mechanism.

## Methods

### Bacterial strains and media

The *E. coli* S17-1λ*pir* strain was used for cloning purposes, and the *E. coli* BL21 (DE3) strain was used for overexpression and purification of all proteins (Supplemental information Table S2). *E. coli* strains were cultured at 37 °C with shaking (250-275 RPM) in Lysogeny Broth (LB) medium with 40 µg/mL kanamycin, 100 µg/mL ampicillin, and/or 10 µg/mL chloramphenicol when selection was required. *V. harveyi* BB120 was recently reclassified as *Vibrio campbellii* BB120 (a.k.a., ATCC BAA-1116) (35), but for consistency in the literature, we refer to it as *V. harveyi*. BB120 and derivatives were cultured at 30 °C with shaking (250-275 RPM) in Luria Marine (LM) medium with 250 µg/mL kanamycin, 5/10 µg/mL chloramphenicol, and/or 50 µg/mL polymyxin B when selection was required. Plasmids were transformed into electrocompetent *E. coli* S17-1λ*pir* cells and subsequently conjugated into *V. harveyi* strains. *V. harveyi* exconjugants were selected using polymyxin B (50 U/mL).

### Bioluminescence assays

Bacterial cultures were back-diluted to OD_600_ = 0.0005 in 50 mL LM in flasks and grown shaking at 275 RPM. For the standard assay (Fig. 1A), optical density (OD_600_) was measured using a spectrophotometer and a Biotek Cytation3 plate reader was used to measure light units per 200 μL. For the high-throughput assay, cells were diluted 30,000X-fold into 200 μl LM and incubated at 30 °C shaking overnight in the Biotek plate reader with readings every 30 minutes. Bioluminescence per cell was determined by dividing the relative light units (RLU) by the OD_600_. For Figures 1B-D and S2, the plate reader was used to determine bioluminescence and optical density.

### Molecular and chemical methods

PCR was performed using Phusion HF polymerase (New England Biolabs). All restriction enzymes and T4 polynucleotide kinase were purchased from New England Biolabs and used according to the manufacturer’s protocol. Site-directed mutagenesis was performed using Phusion HF polymerase (New England Biolabs) following the QuikChange mutagenesis protocol (Stragene). All oligonucleotides were ordered from Integrated DNA Technologies (IDT). Oligonucleotides used for EMSAs and qPCR are listed in Supplemental Information Table S4. PCR products and plasmids were sequenced using Eurofins Genomics. All plasmids used in this study are listed in Supporting Information Table S3, and cloning procedures are available upon request. DNA samples were resolved using 1% agarose (1x TBE). *V. harveyi* autoinducer 1 (HAI-1) was synthesized using a previously published protocol (36).

### Construction of deletion/epitope-tagged strains

All *V. harveyi* BB120 and KM669 derivative strains in this study were constructed following a previously published technique (7). Briefly, the pRE112 suicide vector was used to construct unmarked deletions or insertions of epitopes in which 1000 bp of upstream and downstream flanking sequence was cloned into pRE112. The pRE112 derivatives were conjugated into *V. harveyi* and selected on chloramphenicol to induce chromosomal recombination of the plasmid. Subsequently, the plasmid was excised via counterselection on 20-25% sucrose. Cells in which the plasmid excision yielded a non-WT locus were detected via colony PCR. All gene deletions were confirmed by DNA sequencing.

### RNA-seq

To isolate RNA, cells were back-diluted to OD_600_ = 0.005 and grown shaking at 30°C until OD_600_ = 1.0. RNA was collected, extracted, DNase-treated, and purified as described previously (7). RNA-seq was performed as described previously (7). Sequence data were deposited in the National Center for Biotechnology Information Gene Expression Omnibus database (NCBI GEO).

### Chromatin immunoprecipitation

Cells were grown to an OD_600_ of 1.0 and protein/nucleic acids were crosslinked using 1% formaldehyde at 30 °C for 30 minutes. Crosslinking was quenched with the addition of 0.42 M glycine and incubated at 30 °C for 5 minutes. 10 ODs of cells were harvested for each replicate and cell pellets were stored at −80 °C. Cells were lysed using lysis buffer (1x Bugbuster (Milipore), 1% Triton X-100, 1 mM PMSF, 50 µg/mL lysozyme, and 1x protease inhibitors) at room temperature for 30 minutes on a nutator. Protease inhibitor cocktail (100x stock) contained the following: 0.07 mg/mL phosphoramidon (Santa Cruz), 0.006 mg/mL bestatin (MPbiomedicals/Fisher Scientific), 1.67 mg/mL AEBSF (DOT Scientific), 0.07 mg/mL pepstatin A (Gold Bio), 0.07 mg/mL E64 (Gold Bio). For ChIP-qPCR experiments, cell lysates were sonicated using a Branson SFX 150 sonicator (30% amplitude, 10” pulse, 30” rest on ice, 4 cycles). For ChIP-seq experiments, cell lysates were sonicated using a Covaris S220 sonicator (175 W peak incident power, 10% duty factor, 200 cycles per burst, 15-minute treatment time, 7.5 °C). After sonication, lysates were clarified via centrifugation at 15,000 RPM for 15 minutes at 4 °C. FLAG-tagged proteins (LuxR was tagged N-terminally and H-NS was tagged C-terminally) were captured from clarified lysates using anti-FLAG agarose M2 beads (Sigma). Lysates were incubated with the anti-FLAG resin for 90 minutes at room temperature in immunoprecipitation buffer (50 mM HEPES-NaOH pH 7.5, 150 mM NaCl, 1 mM EDTA, 1% Triton X-100, 1 mM PMSF) on a nutator. After incubation with lysate, the beads were washed with immunoprecipitation buffer, wash buffer 1 (50 mM HEPES-NaOH pH 7.5, 500 mM NaCl, 1 mM EDTA, 1% Triton X-100, 1 mM PMSF, 0.1% SDS), wash buffer 2 (10 mM Tris-HCl pH 8.0, 250 mM LiCl, 1 mM EDTA, 0.5% NP-40, 0.5% deoxycholate), and 10 mM TE. Nucleoprotein complexes were eluted from the beads using elution buffer (50 mM Tris-HCl pH 7.5, 10 mM EDTA, 1% SDS) at 65 °C for 30 minutes. Subsequently, nucleic acids were freed from crosslinked protein via proteinase K digestion (1.8 mg/mL proteinase K, 2 hours at 42 °C, 16 hours at 65 °C). ChIPed DNA was purified using a PCR clean up kit (Qiagen). For input samples (an aliquot of the sonicated, clarified lysate prior to incubation with beads), 1:1 ratio of lysate:10 mM Tris-HCl pH 8.0 and 1.8 mg/mL RNase A was added and incubated at room temperature for 10 minutes. Proteinase K treatment of input DNA was performed as described above. All ChIP-seq experiments were performed using three biological replicates. Immunoprecipitated DNA concentration/quality was determined using an Agilent 2200 TapeStation and the DNA was subsequently stored at −20 °C.

### ChIP-seq and analyses

Chromatin immunoprecipitation for ChIP-seq experiments was performed as described above. Sonication was performed using a Covaris S220 water bath sonicator; manufacturers’ protocol was followed to achieve ∼150 bp DNA fragments. Immunoprecipitated chromatic DNA was subjected to library preparation using NEBNext Ultra II DNA Library kit (NEB cat# E7645S). Briefly, DNA fragments were end repaired and 5’ phosphorylated, and a 3’ end Adenosine-tail was added. The products were ligated to a NEB hairpin adapter which contains an uracil at the loop region. The adapter loop was opened by Uracil-Specific Excision Reagent (NEB) to excise the uracil. The products were cleaned with Agencourt AMPure XP beads (Beckman Coulter cat# A63882) but no size selection was applied. 10 (GSF1697) or 12 (GSF2248) PCR cycles were run using 8-nt barcoded oligos from NEBNext Multiplex Oligos barcode kit (NEB cat# 6609S). The libraries were purified with Agencourt AMPure XP beads, multiplexed and sequenced on NextSeq 500 (Illumina) with NextSeq75 High Output v2.5 kit (Illumina cat# 20024906) to generate paired-end 2×42 bp reads. The read sequences were de-multiplexed using bcl2fastq (software versions 2.1.0.31). ChIP-seq datasets were trimmed for quality and adapters using Trimmomatic (v0.33 for all samples except the H-NS Δ*luxR* ChIP H-NS samples which used v0.38) with the following parameters: ILLUMINACLIP:adapters.fa:3:20:6 LEADING:3 TRAILING:3 SLIDINGWINDOW:4:20 MINLEN:30. Reads were mapped to the Vibrio harveyi ATCC BAA-1116 genome using bwa (v0.7.17). Peaks were identified using macs2 (v2.1.2) call peak algorithm. To compare H-NS peak heights in the wildtype and ΔluxR mutant backgrounds we had to normalize the data. Naively, we would just normalize each sample based on the number of ChIP-seq reads mapped to the genome but it was unclear whether this would be accurate given the potential interaction between H-NS and luxR. To validate this approach, we identified 98 H-NS peaks that were not associated with luxR binding sites. The coverage underlying the peak in each replicate was normalized to the number of reads mapped to the genome. The mean, calculated across replicates, normalized peak coverage for these 98 luxR independent peaks in the wildtype and ΔluxR strains was plotted. Linear regression shows that there is a significant correlation between these points and that these points generates a line with a slope close to 1 that passes close to the origin. Based on this result we proceeded to normalize H-NS peak heights after normalizing by mapped reads. Distances between LuxR/H-NS and H-NS/H-NS peaks were performed using a custom script. Points were plotted as a function of distance from LuxR peak to the nearest H-NS peak (x-axis) against distance from nearest H-NS peak to the next closest H-NS peak (i.e. doublet, y-axis). H-NS doublet peaks were identified using a custom script that searched for H-NS peaks within 2 kb of each other that had a trough ≤80% of the lower of the two peaks. Sequence data were deposited in the NCBI GEO database.

### Quantitative PCR

ChIP input and elution samples were diluted 1:100 and 1:4, respectively, using nuclease free H_2_O. qPCR reactions were performed using SensiFAST SYBR Hi-ROX (Bioline) according to the manufacturer’s guidelines. qPCR primers were designed to fit the following parameters: amplicon size: 100-105 bp, primer size: 18-22 nt, and primer melting temperature: 56-60 °C. Each reaction contained 2 µL of diluted DNA template (input or elution) and 0.4 µM of each primer (total volume was 10 µL). qPCR reactions were incubated and monitored using a LightCycler 4800 II (Roche). For quantification of relative enrichment, the following equation was utilized: 2^CTinput – CTelution^. All ChIP-qPCR experiments were performed with three biological replicates.

### Purification of LuxR, H-NS, and RNAP proteins

Recombinant, untagged LuxR and *E. coli* 6xHis-RNAP proteins were purified using protocols previously reported (7, 9). H-NS was purified by cloning the *hns* ORF into the pET28B expression vector (Novagen) to generate a 6xHis tag on the C-terminus. The pET28B-*hns*-6xHis plasmid (pRC049) was transformed into NiCo21 *E. coli* (New England Biolabs), which contains lacks several genes encoding proteins that readily contaminate immobilized metal affinity chromatography purifications. The resulting strain, RRC226 (NiCo21-pRC049), was grown at 30 °C to an OD_600_=0.6-0.8 and *hns* overexpression was induced using 1 mM IPTG for 3 hours. Induced cell pellets were resuspended in lysis buffer (20 mM Tris-HCl pH 8.0, 300 mM NaCl, 10 mM imidazole, 1x protease inhibitors, 1 mM PMSF, 0.2 mg/mL DNaseI (GoldBio), 1× FastBreak (Millipore)) and incubated at room temperature for 35 minutes. The protease inhibitor mix included the following: 0.07 mg/mL phosphoramidon (Santa Cruz), 1.67 mg/mL AEBSF (DOT Scientific), 0.07 mg/mL pepstatin A (DOT Scientific), 0.07 mg/mL E-64 (Gold Bio), and 0.06 mg/mL bestatin (MPbiomedicals/Fisher). Clarified lysate was loaded onto a 5 mL His-Trap Ni-NTA column (GE Healthcare Life Sciences) using an Akta Pure (GE Healthcare Life Sciences). Protein was eluted from the column using a linear gradient of elution buffer (20 mM Tris-HCl pH 8.0, 300 mM NaCl, 1 M imidazole). Fractions were analyzed by SDS-PAGE to confirm the presence of 6xHis-H-NS and pooled together. Pooled fractions were concentrated using 10 kDa cutoff centrifugal filters (Sartorius) and dialyzed overnight in against storage buffer (20 mM Tris-HCl pH 8.0, 300 mM NaCl, 1 mM EDTA, 5% glycerol). Dialyzed protein was aliquoted, snap frozen in liquid N_2_, and stored at −80 °C.

### Electrophoretic mobility shift assays

PCR was used to produce promoter substrates; P_*luxC*_, P_02100_, and P_03626_ were amplified using RC003/042, RC549/606, and RC604/605, respectively. Binding reactions were performed using either 1 nM radiolabeled DNA or 8.5 nM non-labelled DNA in EMSA binding buffer (50 mM HEPES-NaOH pH 7.5, 50 mM NaCl, 5% glycerol). Poly(dI-dC) (10 ng/µL, Sigma) was used as non-specific competitor DNA when noted. Reactions were incubated at room temperature for 15 minutes. For competitive binding experiments, reactions were supplemented with the secondary protein in binding buffer and incubated at room temperature for an additional 15 minutes. Reactions containing small substrates were resolved on 6% polyacrylamide (19:1) 1x TBE (100 mM Tris, 100 mM boric acid, 2 mM EDTA) gels using 25 mA constant current for 30-40 minutes at room temperature. These gels were dried down and visualized using a phosphor storage screen (imaged with a GE Life Technologies Typhoon 9500). Reactions containing large substrates were resolved on 6% polyacrylamide (37.5:1) 1x TBE gels using 20 mA constant current for 50-70 minutes at 4 °C. DNA was detected by staining the gel with 1 µg/mL ethidium bromide for 20 minutes at room temperature. To detect displaced proteins in competitive EMSAs, the ethidium bromide-stained gel was transferred to a nitrocellulose membrane. Proteins were detected using anti-His-HRP (Sigma) and anti-FLAG-HRP (Sigma) antibodies.

## Acknowledgements

The authors thank Chelsea Simpson, Evan Schneider, and Victoria Lydick for excellent technical assistance. We thank Jun Liu at the Indiana University Center for Bioinformatics and Genomics for assistance with ChIP-seq and RNA-seq processing and analyses.

## Supplemental Information

**Figure S1.**
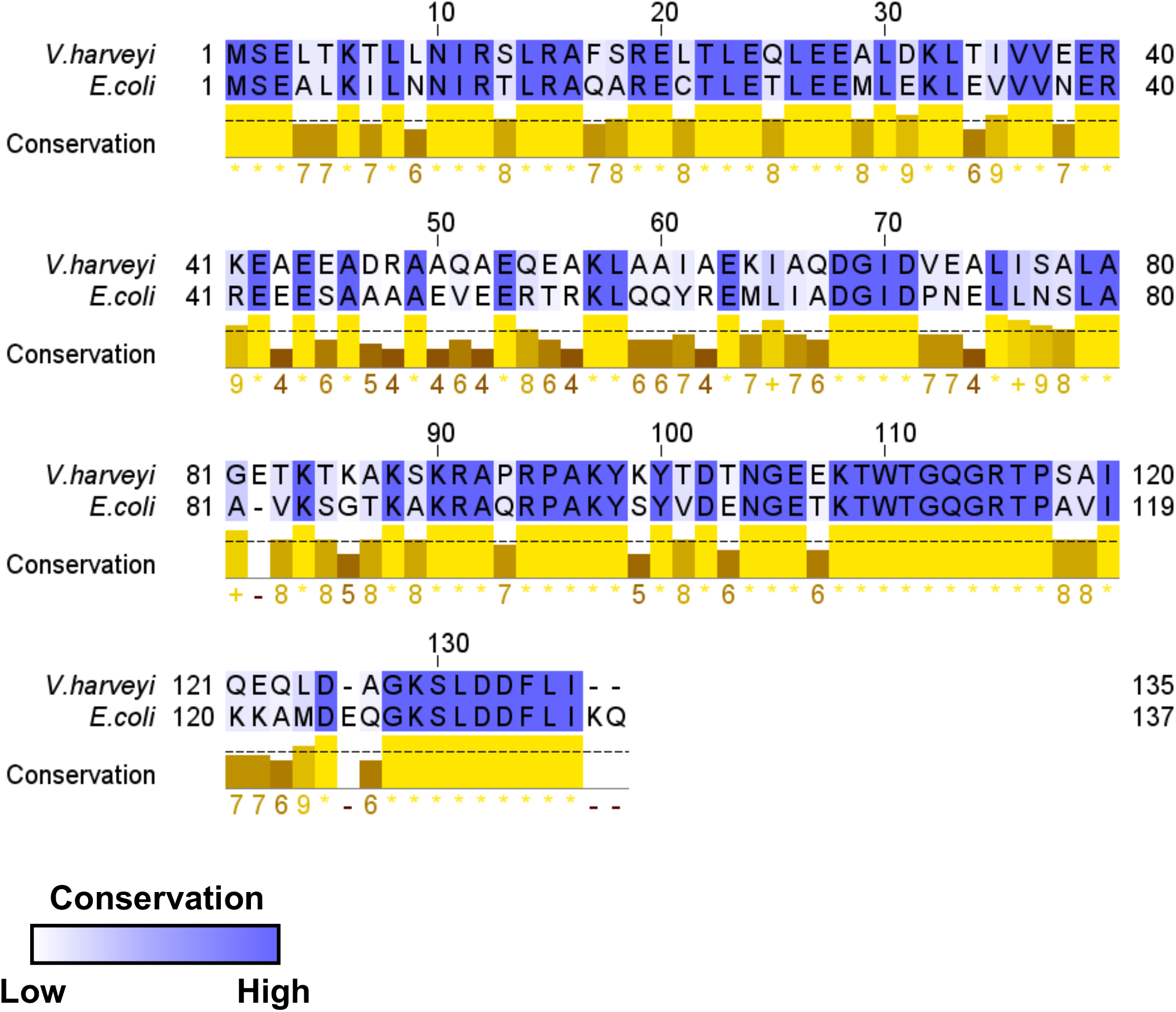
Amino acid identity between *V. harveyi* H-NS and *E. coli* H-NS. The protein sequences for *E. coli* H-NS (accession AYG19681) and *V. harveyi* H-NS (VIBHAR_01827; accession ABU70796) were aligned using Clustal Omega (Madeira *et al*. (2019) *Nucleic Acids Research*) and the diagram generated using Jalview 2.0 (Waterhouse *et al.* (2009) *Bioinformatics*).

**Figure S2.**
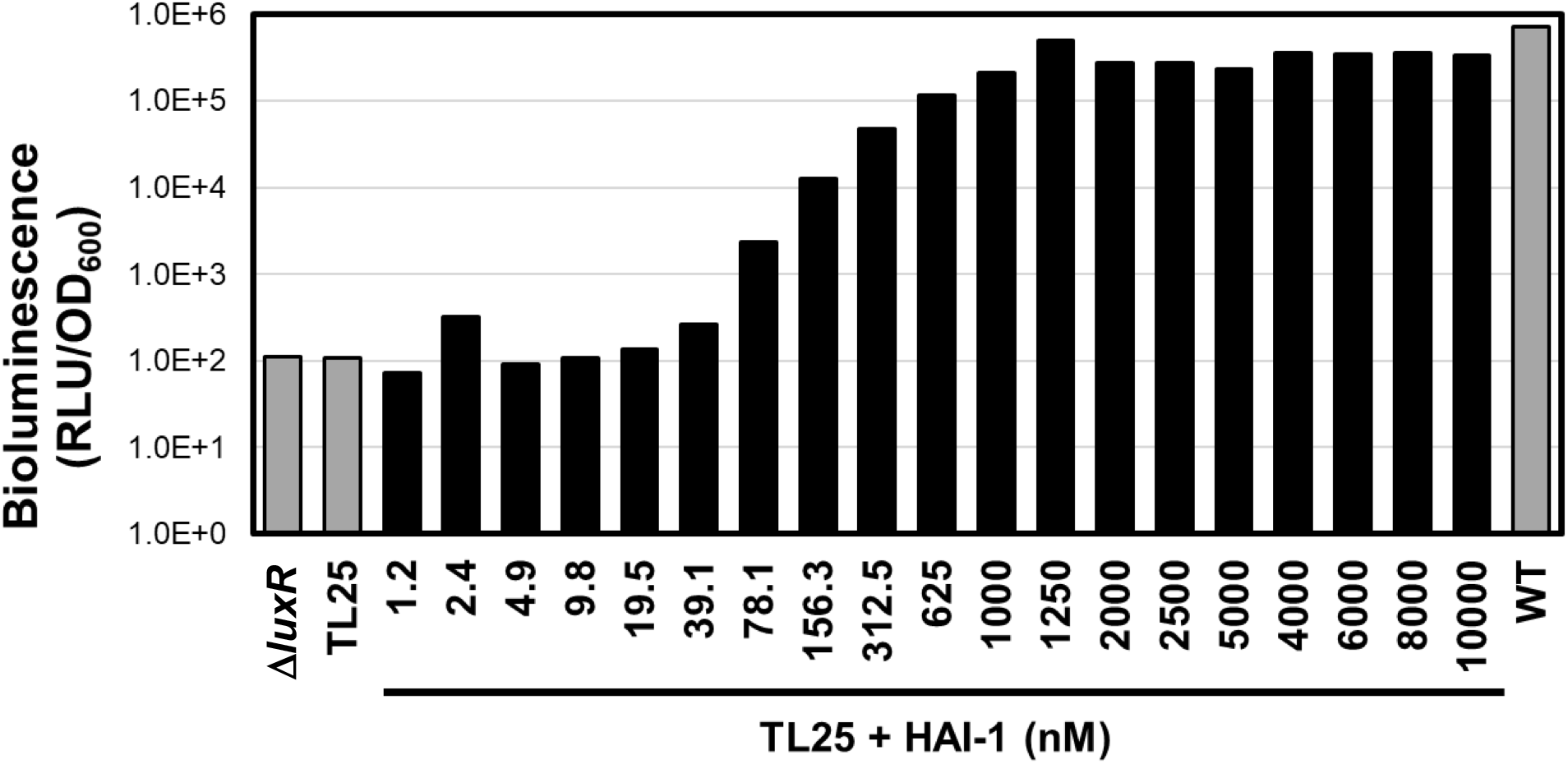
Bioluminescence production in response to AI-1 concentration. Bioluminescence (relative light units (RLU) divided by OD_600_) was measured from wild-type BB120 (WT), Δ*luxR* (KM669), or Δ*luxPQ* Δ*luxM* Δ*cqsS* (TL25) in the presence (black bars) or absence (gray bars) of HAI-1 at the specified concentrations.

**Figure S3.**
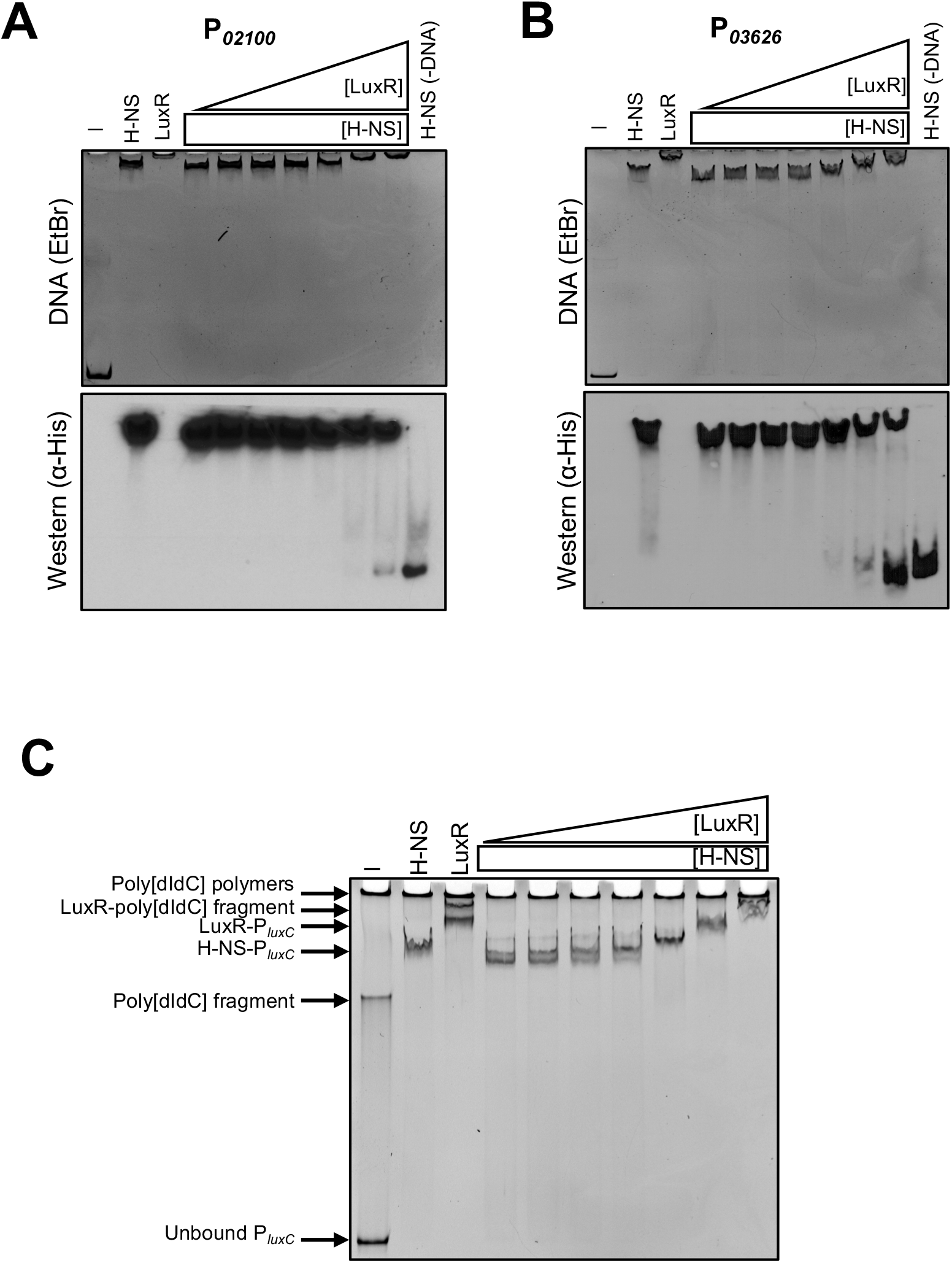
Competition between LuxR and H-NS at promoters. (A) Competitive EMSA reactions containing 9.5 nM P_02100_ DNA (−396 bp from ATG) and 300 nM 6xHis-H-NS and either 31.25, 62.5, 125, 250, 500, 1000, or 2000 nM LuxR. Lanes labelled ‘H-NS’, ‘LuxR’, ‘-’, and ‘H-NS (−DNA)’ contained 300 nM 6xHis-H-NS, 2000 nM LuxR, no protein, and 300 nM H-NS without DNA, respectively. (B) Same experimental set up as in (A) but reactions contained 8.8 nM P_03626_ DNA (−427 bp from ATG) and 200 nM 6xHis-H-NS. All reactions were run on polyacrylamide gels, stained with ethidium bromide (top), and transferred to nitrocellulose membranes and western blot analyses performed (bottom); α-His-HRP antibodies were used to probe 6xHis-H-NS. (C) Same experimental set up from (A) but reactions contained 150 ng poly dIdC, 600 nM 6xHis-H-NS, and P_*luxC*_ DNA.

**Figure S4.**
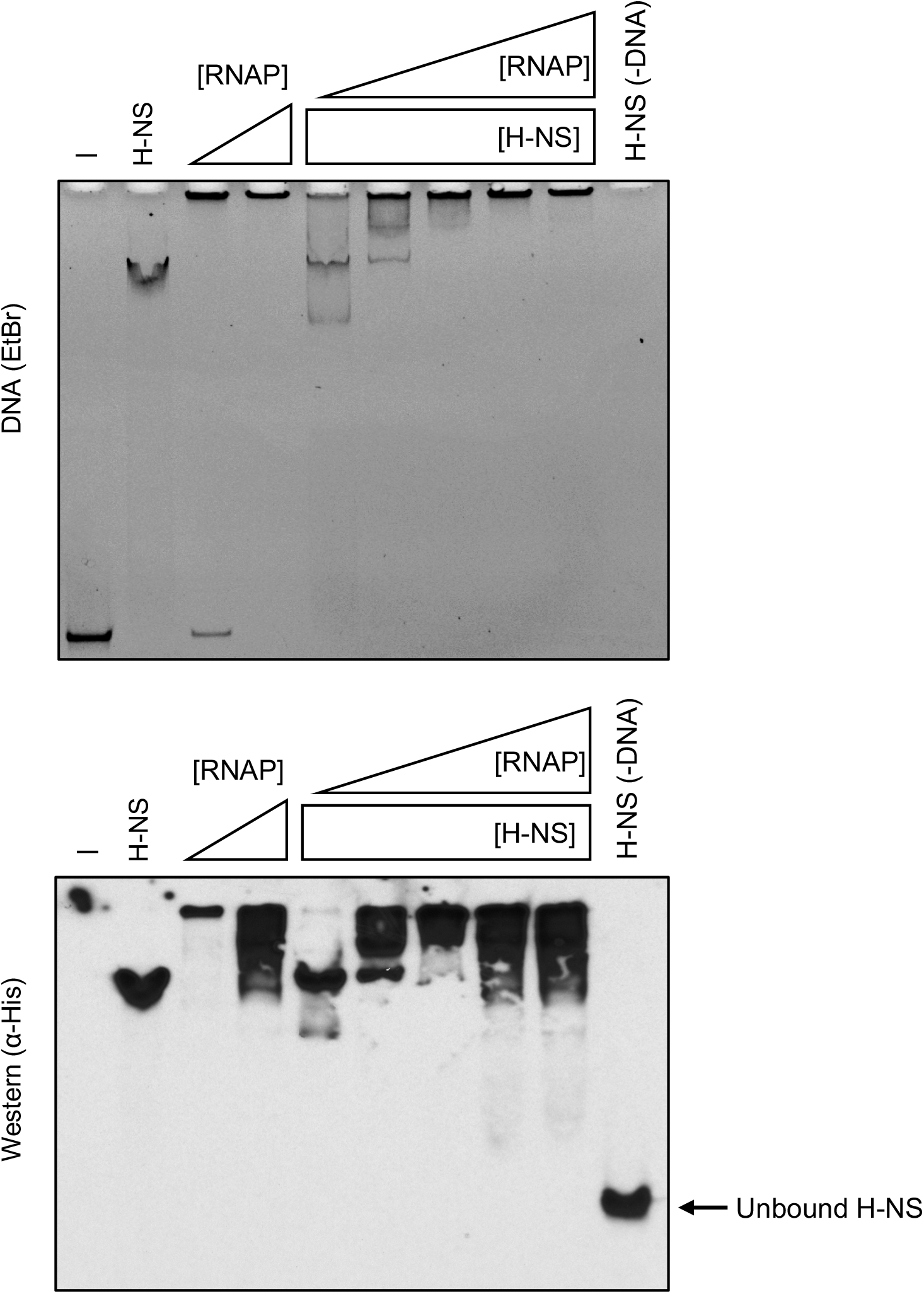
Competition between RNAP and H-NS at P_*luxC*_. (Top) Competitive EMSA reaction containing 8.8 nM Pl_*uxC*_ DNA and 200 nM 6xHis-H-NS and either 31.25, 62.5, 125, 250, or 500 nM RNAP. Lanes labelled ‘H-NS’, ‘RNAP’, ‘-’, and ‘H-NS (−DNA)’ contained 200 nM 6xHis-H-NS, 150/500 nM RNAP, no protein, and 200 nM H-NS without DNA, respectively. Reactions were run on a polyacrylamide gel and stained with ethidium bromide. (Bottom) Reactions from (top) were transferred to a nitrocellulose membrane and probed for 6xHis-H-NS using α-His-HRP antibodies.

**Figure S5.**
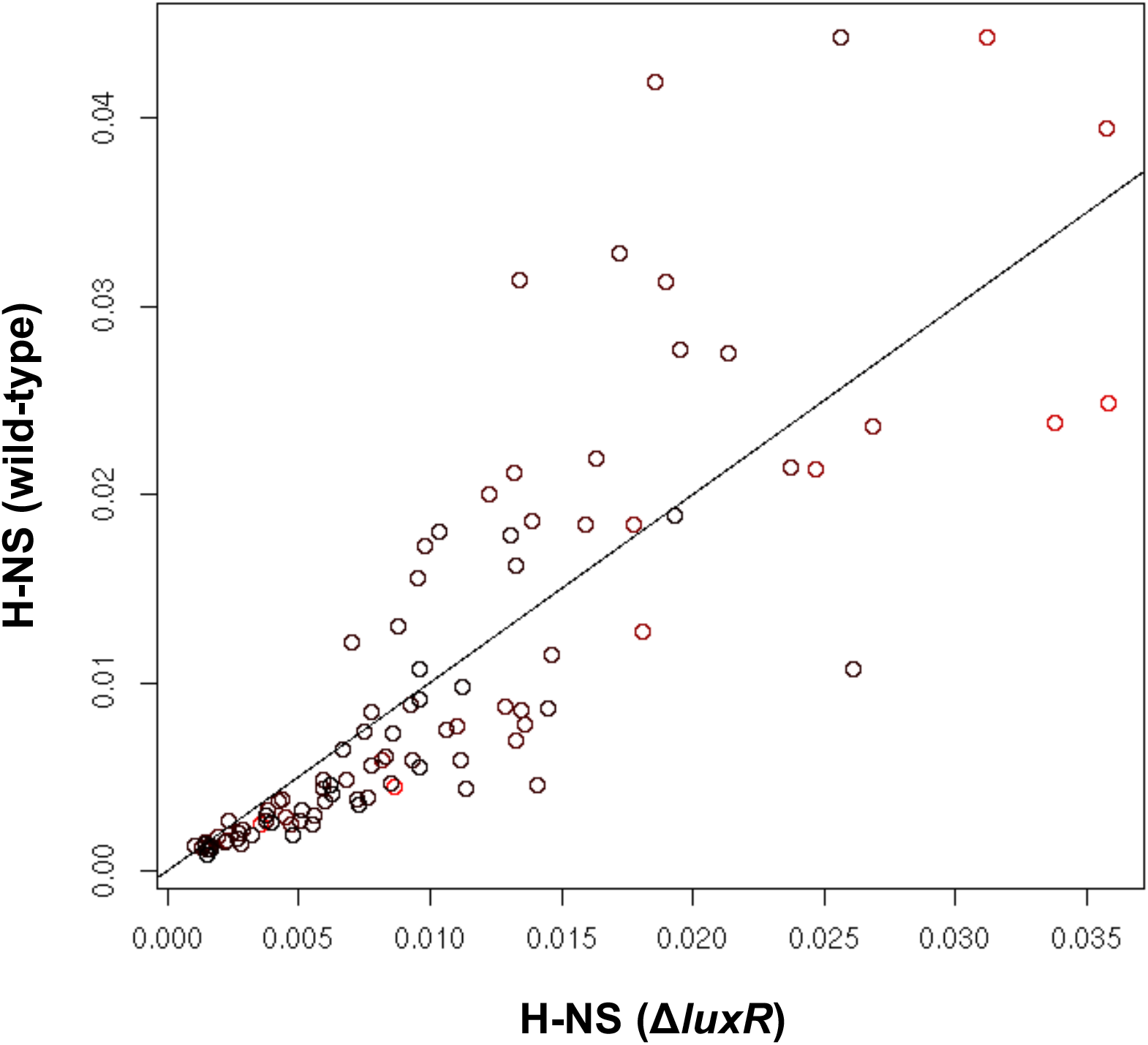
Normalization of H-NS ChIP-seq reads. 98 LuxR-independent H-NS loci were normalized by number of reads mapped per sample and window size. Mean data for each peak in each sample (wild-type vs. Δ*luxR*) are plotted. Data points are colored from black to red to indicate the abundance of proximal LuxR. The correlation between the two data sets is 0.8296.

**Table S1.**
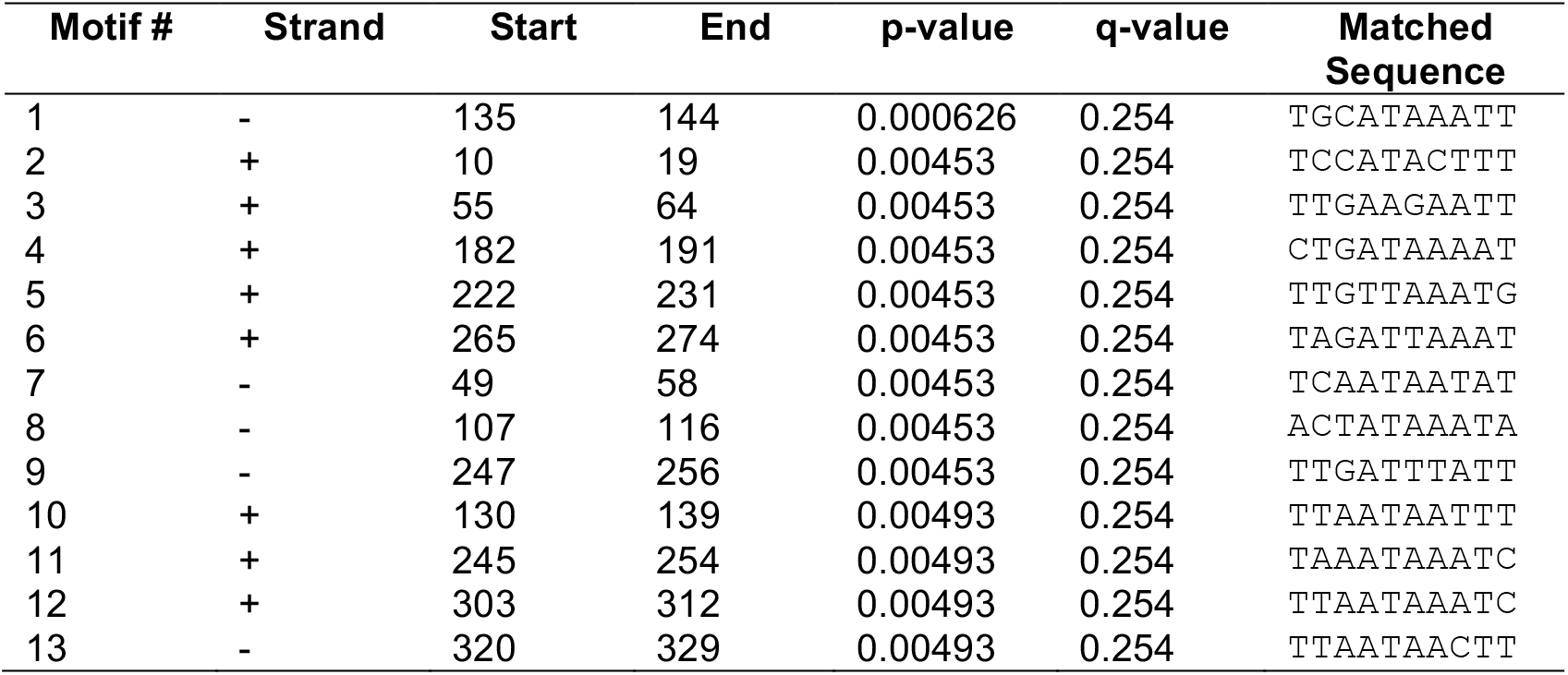
H-NS motifs in P*luxC* using FIMO.

**Table S2.**
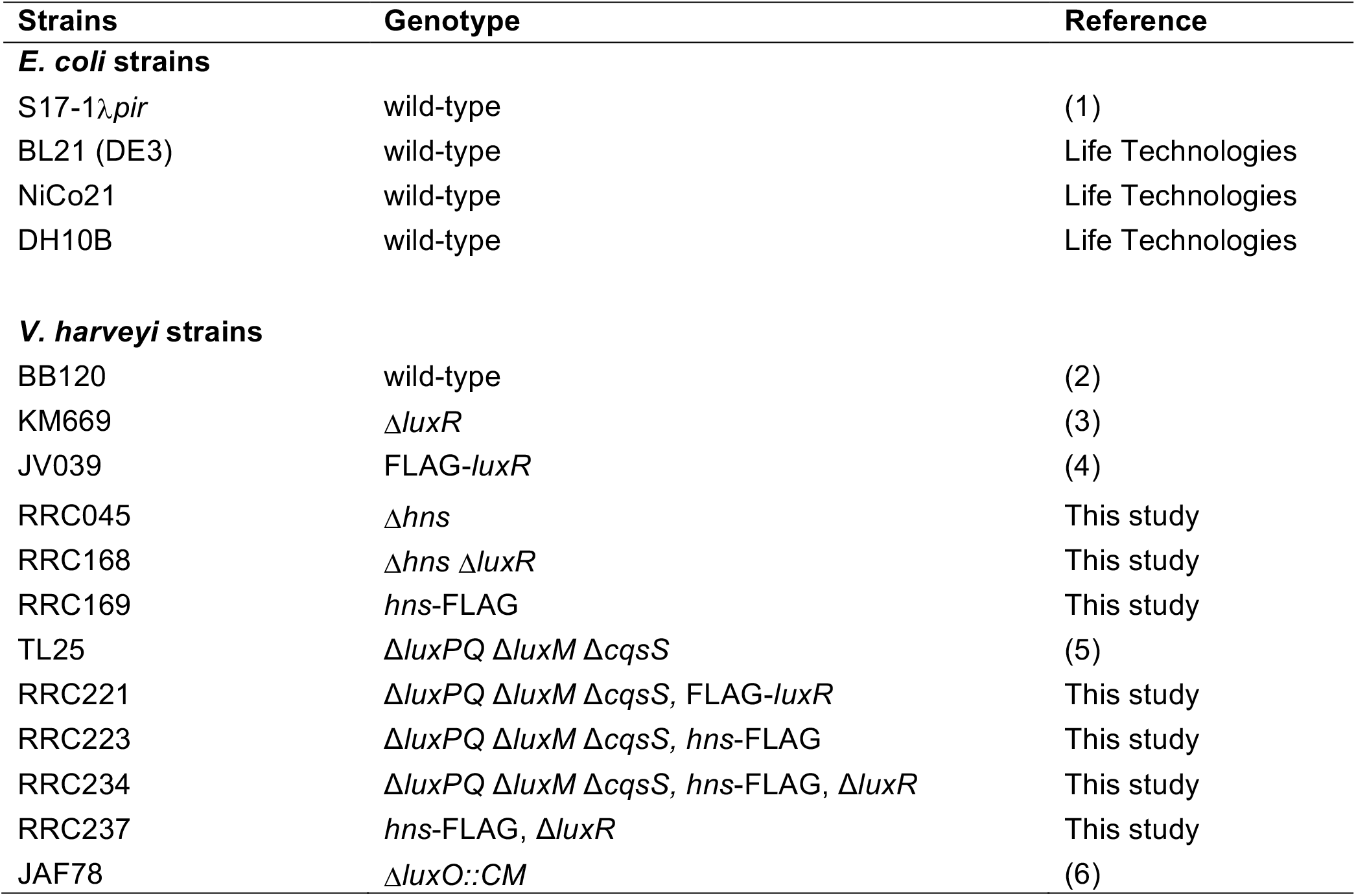
Strains used in this study.

| Strains | Genotype | Reference |
| --- | --- | --- |
| ***E. coli* strains** |  |  |
| S17-1λ*pir* | wild-type | (1) |
| BL21 (DE3) | wild-type | Life Technologies |
| NiCo21 | wild-type | Life Technologies |
| DH10B | wild-type | Life Technologies |
| ***V. harveyi* strains** |  |  |
| BB120 | wild-type | (2) |
| KM669 | $\Delta luxR$ | (3) |
| JV039 | FLAG-*luxR* | (4) |
| RRC045 | $\Delta hns$ | This study |
| RRC168 | $\Delta hns \Delta luxR$ | This study |
| RRC169 | *hns*-FLAG | This study |
| TL25 | $\Delta luxPQ \Delta luxM \Delta cqsS$ | (5) |
| RRC221 | $\Delta luxPQ \Delta luxM \Delta cqsS$, FLAG-*luxR* | This study |
| RRC223 | $\Delta luxPQ \Delta luxM \Delta cqsS$, *hns*-FLAG | This study |
| RRC234 | $\Delta luxPQ \Delta luxM \Delta cqsS$, *hns*-FLAG, $\Delta luxR$ | This study |
| RRC237 | *hns*-FLAG, $\Delta luxR$ | This study |
| JAF78 | $\Delta luxO::CM$ | (6) |

**Table S3.** Plasmids used in this study.

| **Name** | **Description** | **Reference** |
| --- | --- | --- |
| pRE112 | gene deletion vector, *cm<sup>R</sup>* (chloramphenicol resistant), *sacB* (sucrose intolerant), R6Kg origin | (7) |
| pMMB67EH-kanR | *P<sub>tac</sub>* promoter empty vector, IncQ origin, *kan<sup>R</sup>* (kanamycin resistant) | (Simpson *et al.*, submitted) |
| pRC009 | *hns* deletion construct in pRE112, *cm<sup>R</sup>* | This study |
| pJV066 | FLAG-luxR allelic replacement construct in pLAFR2, *cm<sup>R</sup>* | (4) |
| pRC034 | *hns*-FLAG allelic replacement construct in pRE112, *cm<sup>R</sup>* | This study |
| pRC053 | *P<sub>tac</sub>-hns* (*V. harveyi*) complementation vector in pMMB67EH-kanR, *kan<sup>R</sup>* | This study |
| pRC054 | *P<sub>tac</sub>-hns* (*E. coli*) complementation vector in pMMB67EH-kanR, *kan<sup>R</sup>* | This study |
| pRC055 | *P<sub>tac</sub>-hns<sup>L30P</sup>* (*V. harveyi*) complementation vector in pMMB67EH-kanR, *kan<sup>R</sup>* | This study |
| pRC049 | *hns*-6xHis overexpression plasmid in pET28b, *kan<sup>R</sup>* | This study |
| pRC050 | *luxR* deletion construct in pRE112, *cm<sup>R</sup>* | This study |
| pAI900 | a, b, b', and $\omega$ subunits with a C-terminal H10 His tag on b', *amp<sup>R</sup>* | (8), Gourse Lab unpublished [see (9) for construction of similar vectors] |

**Table S4.**
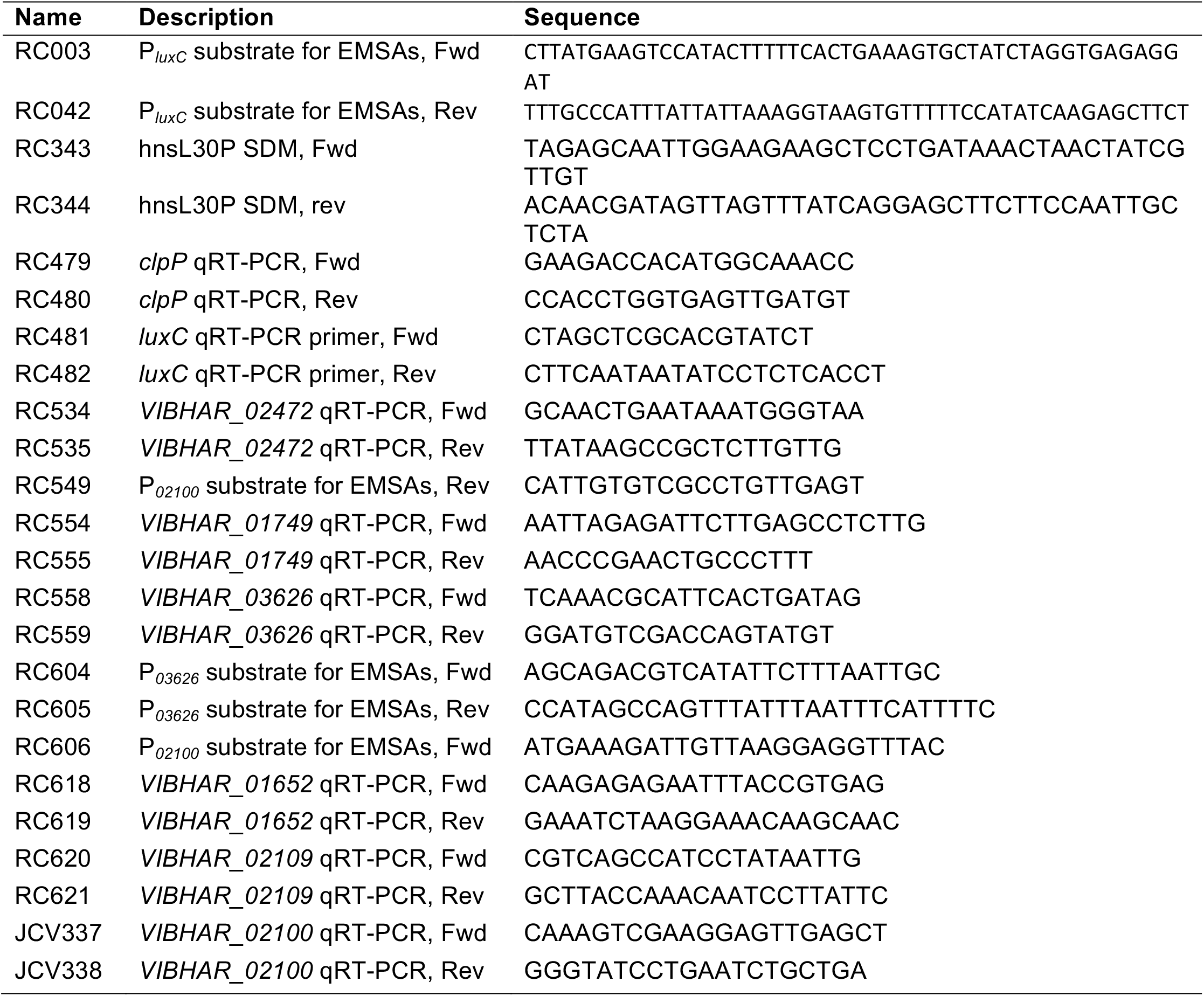
Oligonucleotides used in this study.

